# Fast, Rate-Independent, Finite Element Implementation of a 3D Constrained Mixture Model of Soft Tissue Growth and Remodeling

**DOI:** 10.1101/2020.02.27.968768

**Authors:** Marcos Latorre, Jay D. Humphrey

## Abstract

Constrained mixture models of soft tissue growth and remodeling can simulate many evolving conditions in health as well as in disease and its treatment, but they can be computationally expensive. In this paper, we derive a new fast, robust finite element implementation based on a concept of mechanobiological equilibrium that yields fully resolved solutions and allows computation of quasi-equilibrated evolutions when imposed perturbations are slow relative to the adaptive process. We demonstrate quadratic convergence and verify the model via comparisons with semi-analytical solutions for arterial mechanics. We further examine the enlargement of aortic aneurysms for which we identify new mechanobiological insights into factors that affect the nearby non-aneurysmal segment as it responds to the changing mechanics within the diseased segment. Because this new 3D approach can be implemented within many existing finite element solvers, constrained mixture models of growth and remodeling can now be used more widely.

## 1. Introduction

A distinguishing feature of soft biological tissues is their ability to grow (change mass) and remodel (change microstructure) in response to diverse stimuli, often mechanical and chemical. Multiple approaches for mathematically modeling such growth and remodeling (G&R) have proven useful in describing diverse situations for many different tissues [1–5]. Among these approaches, a constrained mixture model has proven particularly useful when there is a need to account for the different natural configurations, material properties, and rates of turnover of the individual constituents that define the tissue [6]. The classical (heredity integral-based) formulation of this mixture approach is computationally expensive, however, hence most implementations have focused on simple geometries (e.g., cylindrical bodies or axisymmetric membranes).

Herein, we exploit a recent concept of mechanobiologically equilibrated G&R [7] and show computational advantages for illustrative cases that allow direct comparison with full constrained mixture solutions. We also extend prior kinematics to account for general three-dimensional G&R with finite deformations and possible rotations. The proposed rate-independent framework can compute evolving homeostatic states efficiently by enforcing mechanical and mechanobiological equilibrium without the need to track the past history of deposition and removal, as in integral-based approaches, or to integrate evolution equations, as in rate-based approaches. We submit that this new 3D formulation, which can be implemented easily within existing finite element solvers though with a non-symmetric tangent stiffness matrix, will enable fast, reliable finite element simulations of many G&R problems while accounting for critical differences in the diverse constituents that characterize soft tissues.

## 2. A mechanobiologically equilibrated constrained mixture model

We first review local equations for mechanobiologically equilibrated mass fractions, deformation gradients, strain energy functions, and stresses, at constituent and mixture levels, which when complemented with an equilibrium value for a given stimulus function for mass production furnish a set of equations to compute fully resolved states at any material point and G&R time *s.*

### 2.1. Mechanobiological equilibrium

Consider an in vivo loaded configuration *κ* of a soft tissue that consists of a mixture of multiple constituents, in particular, various types of cells, extracellular matrix proteins, glycosaminoglycans, and abundant water. Because effects of internal solid-fluid interactions are negligible with respect to other characteristics exhibited by these tissues for applications of interest (e.g., ex vivo testing and in vivo behaviors during cyclic loading), we do not account explicitly for a solid-fluid mixture with momentum exchanges. Rather, the tissue is modeled as a constrained mixture of multiple hydrated solid constituents, with the hydrostatic pressure associated with interstitial water absorbed by a Lagrange multiplier that enforces transient incompressibility. In a continuum theory of constrained mixtures for G&R [6], properties at a material point in configuration *κ* are represented in a locally averaged sense, in terms of multiple structurally significant constituents *α* = 1, …, *N*, to satisfy mass balance in spatial form

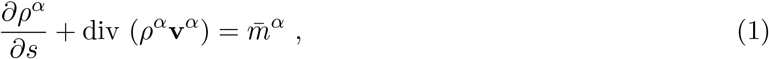

where *ρ^α^* is the homogenized, apparent mass density (mass of constituent *α* per unit current volume of mixture), **v**^*α*^ the velocity (*constrained* to equal the velocity **v** of the mixture), and 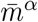 the net rate of mass density production or removal, which must be prescribed constitutively. Let 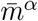 be defined in terms of true rates of mass density production *m^α^* > 0 and removal *n^α^* > 0 as 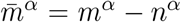, both of which must be prescribed constitutively. It proves useful to define a stimulus function **ϒ**^*α*^ = *m^α^/n^α^* > 0, which enhances (**ϒ**^*α*^ − 1 > 0), reduces (**ϒ**^*α*^ − 1 < 0), or balances (**ϒ**^*α*^ − 1 = 0) mass production with respect to mass removal. Constitutively prescribing *m^α^* and *n^α^* is tantamount to prescribing *n^α^* and **ϒ**^*α*^, which is often more convenient from a modeling perspective. Because div(*ρ^α^***v**^*α*^) = *ρ^α^* div **v**^*α*^ + **v**^*α*^ grad *ρ^α^*, thus *∂ρ^α^/∂s* + **v**^*α*^·grad 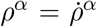, with 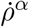 the material time derivative of *ρ^α^*, and 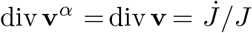, with *J* = det **F** the Jacobian determinant of a deformation gradient **F** defined at the mixture level, which conveniently describes (measurable) deformations between an initial, *original* homeostatic configuration *κ_o_* and *κ*. Equation (1) can thus be written in terms of the referential mass density 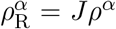 (defined per unit reference volume of mixture) as

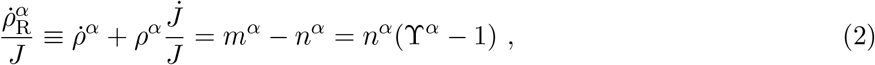

which generalizes rate equations for 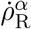 obtained previously [7, 8] from an integral-based approach based on a first-order kinetic decay *n^α^ = k^α^ρ^α^*, with *k^α^* a rate parameter that defines the removal. Similarly, spatial linear momentum balance for constituent *α*, with vanishing interactive forces among constituents due to their constrained motion, may be written as

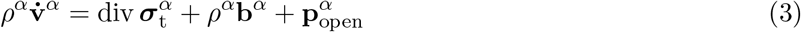

where 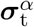 is the total Cauchy stress tensor (i.e., including a contribution ***σ**^α^* derived from a strain energy function and others arising from kinematic constraints, such as incompressibility) for a constituent at the mixture level and **b**^*α*^ is the constituent body force per unit current mass of constituent; additionally, 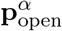 represents the (excess) exchange of momentum not caused by the net exchange in mass 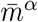, which is typically zero in soft tissues.

Summation of mass (2) and momentum (3) balances over all constituents, with **v**^*α*^ = **v** and 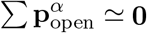, yields the mixture relations

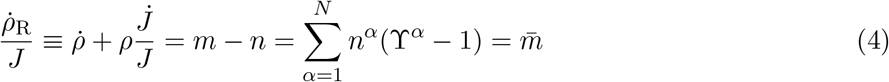

and

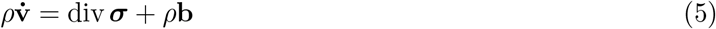

where 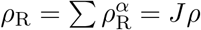, *m* = ∑ *m^α^, n* = ∑ *n^α^*, 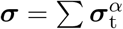, and *ρ***b** = ∑ *ρ^α^***b**^*α*^.

As in [7], and because *n^α^* > 0, we observe from Eqs. (2) and (4) that a sufficient condition for a soft tissue to be in mechanobiological equilibrium (to preserve its mixture mass, composition, and properties) is

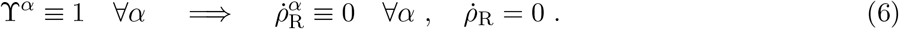

Mechanical (static) equilibrium in a homeostatic state (denoted by subscript *h*) additionally requires, from Eq. (5),

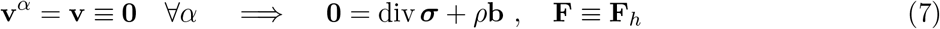

with **F**_*h*_ describing deformations between *κ_o_* and any *homeostatic* configuration *κ_h_*, original or evolved (Fig. 1), which allows adaptive homeostasis [9]. Importantly, Eqs. (6) and (7) also approximate G&R processes wherein the characteristic rate for adaptation is faster than the rate of change of the stimulation, thus yielding mechanobiologically quasi-equilibrated G&R evolutions [10, 11].

**Figure 1:**
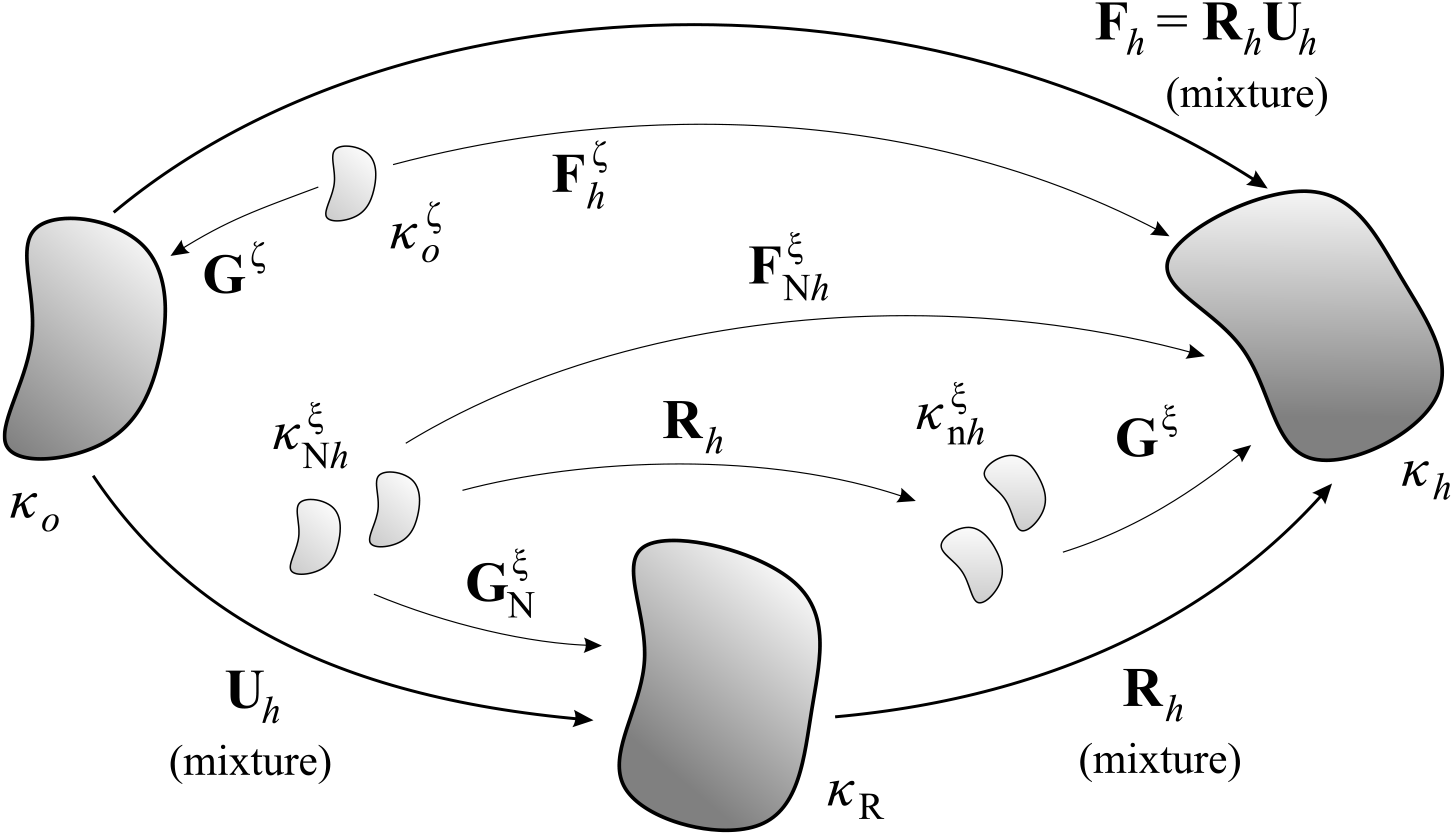
Schematic representation of different configurations *κ_i_* involved in mechanobiologically equilibrated arterial G&R, with *ξ* ≡ collagen (c) and smooth muscle (m) experiencing continuous turnover but *ζ* ≡ elastin (e) not turning over. Note that *κ_o_* and *κ_h_* are both homeostatic; all configurations are in vivo.

### 2.2. Mechanobiologically equilibrated mass fractions

To generalize the present formulation, consider two types of load-bearing constituents *α = ζ* ∪ *ξ* within a soft tissue at G&R time *s* = 0 that evolve differently for *s* > 0. Let constituents *ζ* = 1, .., *N^ζ^* not turnover, whereby their mass remains constant and their response can be described with (rate-independent) hyperelasticity. In contrast, let constituents *ξ* = 1, .., *N^ξ^* turnover continuously within extant matrix, thus contributing to local changes in mass and microstructure, with their response described with rate-independent G&R. Respective initial mass fractions 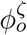 and 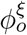 satisfy 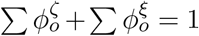, with *N^ζ^ + N^ξ^ = N*. Examples within the arterial wall are functional elastic fibers (= *ζ*), consisting of elastin and associated microfibrils, which are produced during the perinatal period and have a half-life of decades, and collagen fibers (= *ξ*), which are produced continuously and have a half-life of weeks to months. Because differential production and removal of constituents *ξ* contribute to changes in mass of the mixture, both types of constituents can present evolved mass fractions 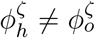 and 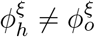 at *κ_h_*, satisfying 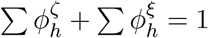, with

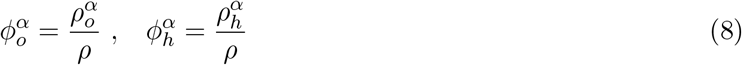

for all constituents *α* = 1, …, *N^ζ^ + N^ξ^*, where 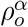 and 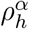 are equilibrated mass densities (defined locally at *o* or *h*, respectively), and *ρ* is the actual mass density of the overall tissue (mixture), which we assume to be constant due to the highly hydrated states of all solid constituents.

#### 2.2.1. Constituents that do not turnover

Because the mass of constituent *ζ* remains constant, a Piola transformation yields 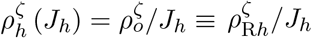, thus

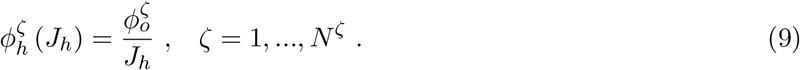

#### 2.2.2. Constituents that turnover

To delineate differential changes in mass of constituents *ξ* under mechanobiological equilibrium, one needs first to describe how they evolve with respect to each other under general G&R. For example, without explicitly prescribing *n_ξ_* and **ϒ**^*ξ*^ in Eq. (2), let all constituents respond to changes in stimuli with proportional out-of-equilibrium stimulus functions and mass-specific rates for removal: 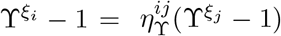 and 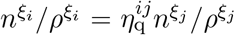 for *ξ_i_* ≠ *ξ_j_* = 1, …, *N^ξ^*, with 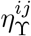 and 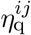 respective proportionality ratios. Then, from Eq. (2), local changes in constituent mass per respective unit mass satisfy

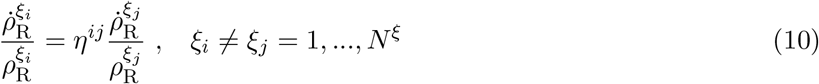

with 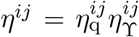. Exact integration of Eqs. (10) from state *κ_o_* (for which 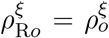) to *κ_h_* yields the following *N^ξ^* − 1 independent relations among the mass fractions 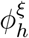

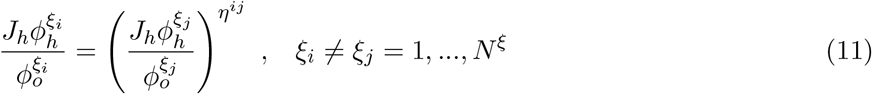

where we used 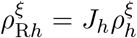 and Eqs. (8), which are equivalent to those obtained in [7] between the two evolving constituents considered therein (smooth muscle cells “*m*” and collagen fibers “*c*”, with a single *η = η_q_η*_ϒ_). Finally, the constraint 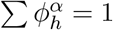, with 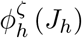 from Eq. (9), requires

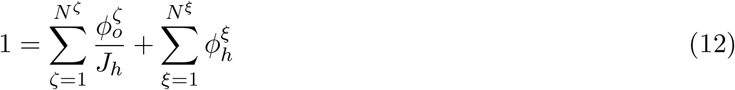

which, along with Eqs. (11), form a system of N independent equations that provide implicit expressions for 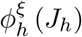. If *η_ij_* = 1 ∀{*ξ_i_, ξ_j_*}, then Eq. (11) reduces to 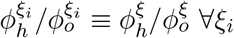, and Eq. (12) to

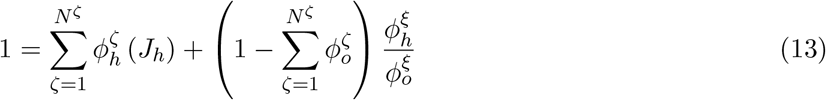

where 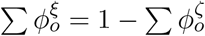, which yields

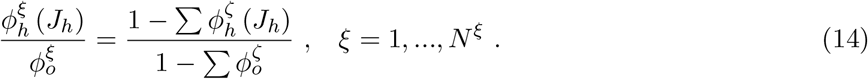

### 2.3. Mechanobiologically equilibrated deformation gradients

In a full constrained mixture theory, constituents are assumed to be deposited within the mixture at deposition time *τ* ≤ *s* in intermediate configurations, relative to their own possibly evolving natural configurations 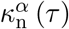, via symmetric and volume-preserving deposition stretch tensors **G**^*α*^ (*τ*). Since the motion of each constituent, once deposited, equals that of the soft tissue, the deformation experienced by the material deposited at time *τ* that survives at *s* is 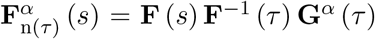 [12]. Because newly deposited constituents at time *τ* satisfy 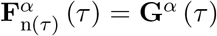, one notices (Appendix A) that 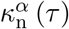 can be interpreted as natural configurations that evolve with the configuration of the mixture *κ*(*τ*), with **G**^*α*^ playing the role of a (spatial) *left* (pre)stretch tensor when referred to a rotated natural configuration 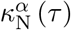 that evolves while attached to the rotated configuration of the mixture *κ*_R_ (*τ*). In that case, it proves convenient to let an associated deformation gradient

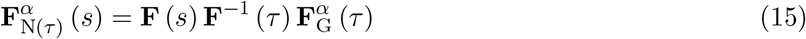

map line elements (fibers) from the rotated natural configuration 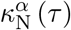 to the current configuration *κ* (*s*), where

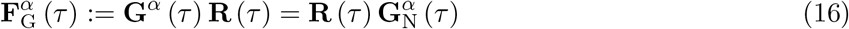

is a constituent-specific deposition tensor at *τ*, with **R** the rotation tensor from a polar decomposition of **F**, thus

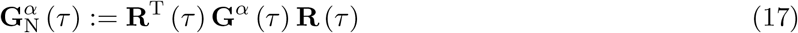

is the associated (symmetric, volume-preserving) *right* (pre)stretch tensor defined in configuration 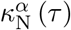 (which is rotated with respect to the spatial configuration 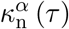 but unrotated with respect to the reference configuration of the mixture *κ*(0)). We then obtain mechanobiologically equilibrated deformation gradients for all constituents *α* from their respective equilibrated natural configurations to the equilibrated current configuration of the mixture *κ_h_*, described by **F** (*s*) = **F**_*h*_ (see Eq. (7) and Fig. 1).

#### 2.3.1. Constituents that do not turnover

Constituents *ζ* are deposited and cross-linked prior to *s* = 0; we account for their elastic response through mechanically equivalent deposition stretch tensors defined at the initial time **G**^*ζ*^ (*τ* = 0) = **G**^*ζ*^ = **const**. Hence, their fictitious natural configurations 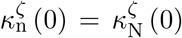 do not evolve over time, but are fixed and attached to the reference configuration for the mixture *κ_o_*. Equation (15) particularized to a mechanobiologically equilibrated state for which **F** (*s*) = **F**_*h*_, with **F** (*τ* = 0) = **I**, yields (see Fig. 1)

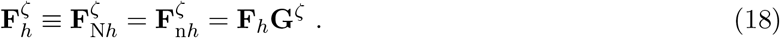

#### 2.3.2. Constituents that turnover

Deposition stretches arise when synthetic cells act on the newly-secreted matrix via actomyosin activity [13], with magnitudes becoming constitutive parameters and so too the orientation of the new tissue when deposited [12, 14]. We assume that the magnitude and orientation of 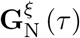 in Eqs. (15)–(17) remain constant ∀_*τ*_, including mechanobiologically equilibrated evolutions ∀_*s*_, thus 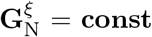. Conversely, 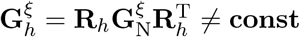, in general, because rotations may arise as the vessel evolves. Equation (15) particularized for constituents that turnover within a homeostatic state, with **F** (*τ*) = **F** (*s*) = **F**_*h*_ ∀_*τ*_, yields (see Fig. 1)

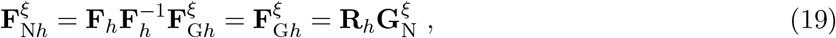

which will prove useful when computing mechanobiologically equilibrated (rotated) Cauchy stresses subject to material frame indifference. Of course, more complex theories can include remodeling of deposition stretches in referential or spatial settings, with either their magnitude (e.g., via a fibrosis-driven maladaptation [15]) or alignment (e.g., via stress-, stretch-, or energy-driven reorientations [12, 16]) evolving over G&R time scales, but we do not consider such cases here for simplicity.

### 2.4. Mechanobiologically equilibrated strain energy

#### 2.4.1. Constituents that do not turnover

Consider strain energy functions 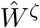 per unit reference volume of their natural configurations 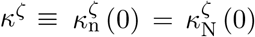. Since constituents *ζ* are deposited within *κ_o_ = κ*(0), associated contributions at the mixture level (per unit reference volume of mixture) are weighted directly by respective *original* volume fractions 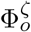 as 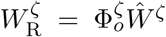. Noting that 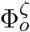 coincides with the mass fraction 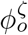 for mixtures of constituents with the same mass density [7], as assumed herein,

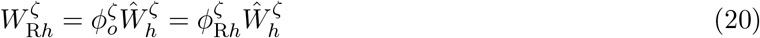

where 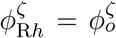 and 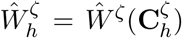 depends on the equilibrated *ζ*-constituent-specific right Cauchy–Green tensor 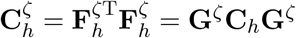, from Eq. (18), with 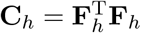 the right Cauchy–Green tensor for the equilibrated mixture.

#### 2.4.2. Constituents that turnover

Consider strain energy functions 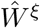 per unit reference volume of their natural configurations 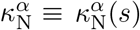, with 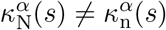 in general. Since constituents *ξ* are deposited within *κ_h_ = κ*(*s*), contributions at the mixture level (per unit current volume of mixture) are weighted by *evolved* volume fractions 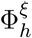 as 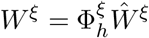, whereupon 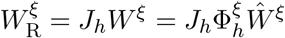. Noting that 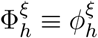,

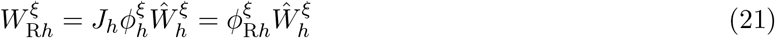

where 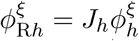 and 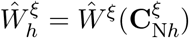 depends on the equilibrated *ξ*-constituent-specific right Cauchy–Green tensor 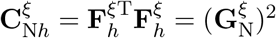, from Eq. (19).

#### 2.4.3. Mixture

Hence, we obtain the following rule-of-mixtures relation for *W*_R*h*_, defined per unit volume in *κ_o_*, in terms of 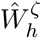, defined per unit volume in *κ^ζ^*, and 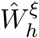, defined per unit volume in 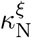

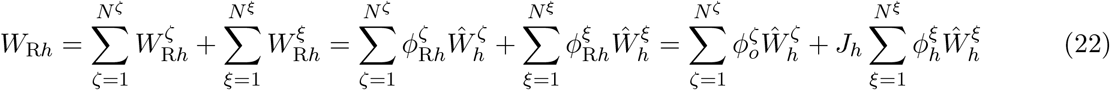

which, importantly, evolves consistent with both limits: 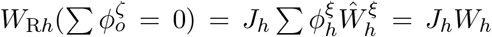 (all constituents turn over) and 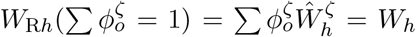 (constituents do not turn over, thereby *J_h_* ≡ 1 relates *W*_R_ and *W*). The mechanobiologically equilibrated strain energy for the mixture reads

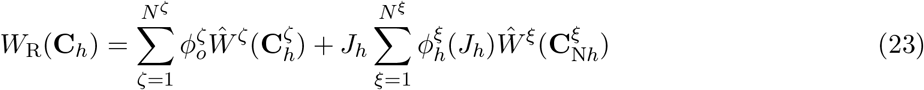

with 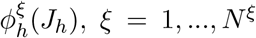, from Section 2.2.2. This energy defines that stored by the mixture as a consequence of its current, pre-stretched homeostatic, in vivo state. An extension of Eq. (23) that allows computation of hyperelastic responses with respect to the configuration *κ_h_* is given in Appendix B.

### 2.5. Mechanobiologically equilibrated stresses

The second Piola–Kirchhoff stress for the mixture **S** during transient loading reads

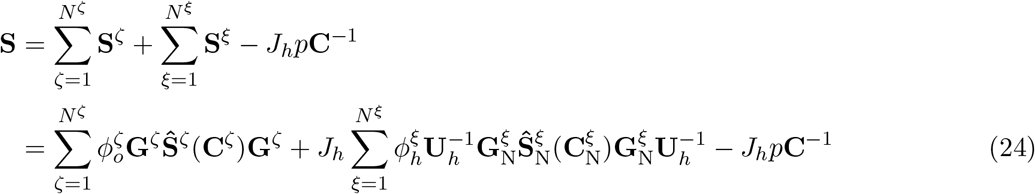

with the appropriate **C, C**^*ζ*^(**C**), and 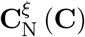 given in Appendix B. Note, too, that the mixture-level Lagrange multiplier *p* associates with the (intermittently imposed) constraint *J = J_h_*, and

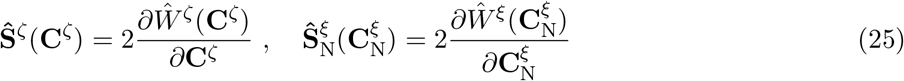

are second Piola–Kirchhoff stresses at the constituent level for both types of constituents. Straightforward particularization of these expressions to **F** = **F**_*h*_ provides mechanobiologically equilibrated stresses at the current in vivo state.

#### 2.5.1. Constituents that do not turnover

**C**^*ζ*^ particularizes to 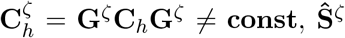 to 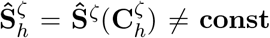 (in general), and **S**^*ζ*^ in Eq. (24) to

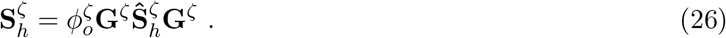

Associated equilibrated Cauchy stresses read

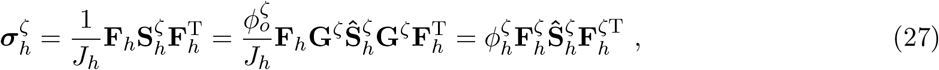

where we used Eq. (18) (see Fig. 1).

#### 2.5.2. Constituents that turnover

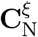 particularizes to 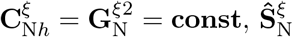 to 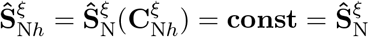, and **S**_*ξ*_ in Eq. (24) to

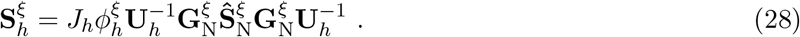

Associated equilibrated Cauchy stresses read

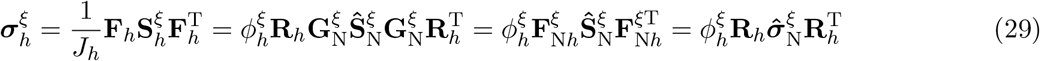

where we used 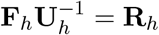, Eq. (19) (see Fig. 1), and a rotated Cauchy stress tensor for constituent ξ at the constituent level

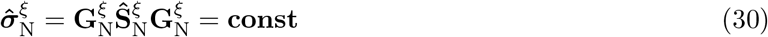

which remains constant during the mechanobiologically equilibrated evolutions considered here.

#### 2.5.3. Mixture

Hence, at homeostatic states **F** = **F**_*h*_, equilibrated second Piola–Kirchhoff stresses for the mixture are

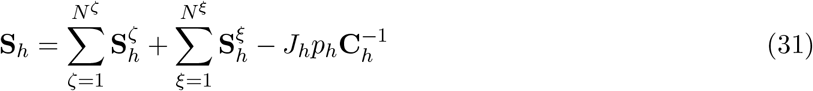

with 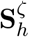 and 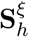 given in Eqs. (26) and (28), respectively. Associated Cauchy stresses, to be used in Eq. (7), specialize to the following evolved rule of mixtures

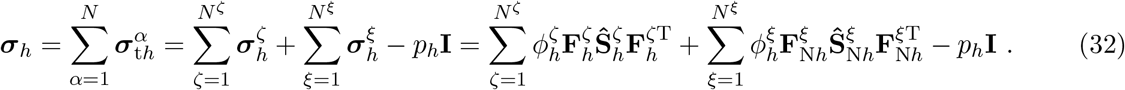

with 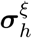 given alternatively in terms of 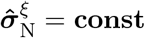 in Eq. (29). Note that 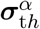 in Eq. (3) includes 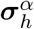, derived from the strain energy, and an associated reaction from the Lagrange multiplier −*p_h_***I**.

### 2.6. Mechanobiologically equilibrated stimulus function and Lagrange multiplier

The hyperelastic stresses 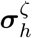 in Eq. (32) are computed from the equilibrated deformation gradient **F**_*h*_, Jacobian-dependent mass fraction 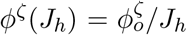, and equilibrated stresses 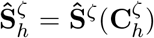, in Eq. (27); in contrast, 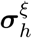, in Eq. (29), are computed from the unique equilibrated rotation **R**_*h*_ from **F**_*h*_, Jacobian-dependent mass fraction *ϕ*^ξ^(*J_h_*), in Section 2.2.2, and the constant tensor 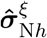, in Eq. (30). Hence, only *p_h_* in Eq. (32) remains unknown, which one obtains by invoking mechanobiological equilibrium conditions in Eq. (6).

Motivated by Fung’s call for mass-and-structure growth-stress relations [17], and consistent with previous constrained mixture models, let *ξ*-constituent-specific stimulus functions *ϒ*^*ξ*^ account for changes in mass production in response to cell-perceived changes in stress relative to homeostatic baseline values. In particular, among other possible equilibrated stimuli, let 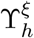 depend on a set of invariants 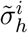 of the total Cauchy stress tensor ***σ**_h_*, and perhaps structural tensors, that include the extent of *p_h_*, that is 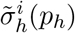 from Eq. (32). Moreover, let the out-of-equilibrium stimulus functions **ϒ**^*ξ*^ − 1 be proportional to each other, as in Section 2.2.2, so equilibrium conditions in Eq. (6) reduce ∀*ξ* to a single nonlinear algebraic equation ϒ_*h*_ = 1. Finally, let 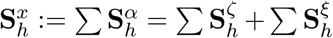 in Eq. (31) be the total “extra” [18], with superscript *x*, second Piola–Kirchhoff stresses, with associated Cauchy stresses 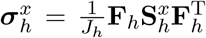 given in Eq. (32), which allows invariants 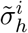 to be expressed in terms of scalar products involving the second-order tensors **C**_*h*_, 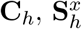, scalars *J_h_, p_h_*, and their couplings, that depend on **C**_*h*_ and *p_h_*. Thus

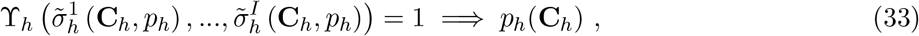

which is a generally implicit relation that yields the (a priori unknown) volumetric contribution to stress at the mixture level in Eq. (32). As an important consequence, the mechanobiologically equilibrated stress field given by Eq. (32), complemented by (33), depends only on the current state of deformation and, hence, is path-independent. Nonetheless, similar to the stress field for a Cauchy elastic material [19], the mechanical work done by this mechanobiologically equilibrated stress field is, in general, path-dependent, which will have further implications regarding its constitutive tangent, as noted in Section 3. The associated material model is summarized in Box 1.

#### Box 1: Mechanobiologically equilibrated constrained mixture model for G&R: kinematics and stresses

i. Deformation gradient for the mixture (from *κ_o_* to *κ_h_*, see Fig. 1) **F**_*h*_ = **R**_*h*_**U**_*h*_, with associated right Cauchy–Green tensor 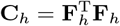
ii. Deformation gradient for constituents *ζ* (from 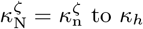), which do not turnover 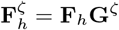, with associated right Cauchy–Green tensor 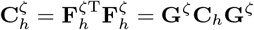
iii. Deformation gradient for constituents *ξ* (from 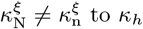), which turnover 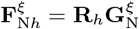, with associated right Cauchy–Green tensors 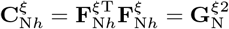
iv. Jacobian (*J_h_* =det **F**_*h*_) dependent mass fractions for different constituents 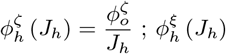 determined implicitly from Eqs. (11) and (12)
v. Second Piola–Kirchhoff stresses for constituents *ζ* (at mixture-level) 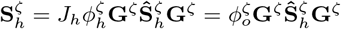, with constituent-level stresses 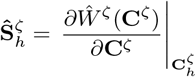
vi. Second Piola–Kirchhoff stresses for constituents *ξ* (at mixture-level) 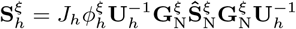, with constituent-level stresses 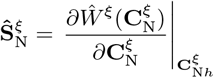
vii. Cauchy stresses for different constituents (at mixture-level) 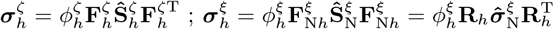 (rotated stresses 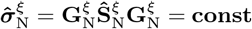)
viii. Consistency parameter (i.e., evolving Lagrange multiplier) enforcing mechanobiological equilibrium 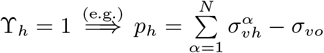, with volumetric stresses 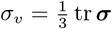
ix. Rule-of-mixtures stresses for the mixture

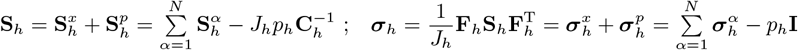

#### Example.

For illustrative purposes and based on previous G&R analyses of prototypical cylindrical arteries [7, 8, 10], consider a linearized 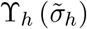, with 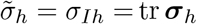 the first principal invariant of ***σ**_h_*, that is,

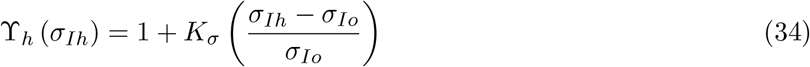

with *K_σ_* a gain parameter for mass production; hence, a possible flow-induced shear stress stimulus for G&R is neglected here for simplicity (see Appendix C for possible accounts of this effect on different 3D computational frameworks). Thus, **ϒ**_*h*_ = 1 yields *σ_Ih_ = σ_Io_*, or, in terms of volumetric (hydrostatic) components of stress 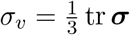,

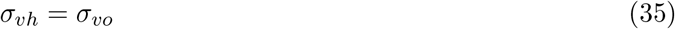

which thereby remains constant during mechanobiologically equilibrated G&R, with its initial value 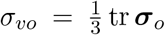 known from the reference state. This choice of stress stimulus, initially motivated to capture the biaxial nature of in vivo wall stresses during arterial G&R adaptations (with the arterial wall subject to transmural pressure and axial stretch), relates closely to other choices in the literature where representative scalars of the volumetric stress contribution were identified as thermodynamic forces driving growth [20, 21]. From Eq. (32), 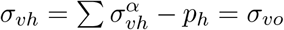, which yields *p_h_* as

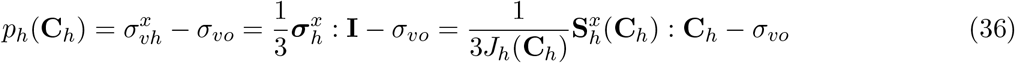

and Cauchy stresses, from Eq. (32), decompose into deformation-dependent deviatoric and constant volumetric components

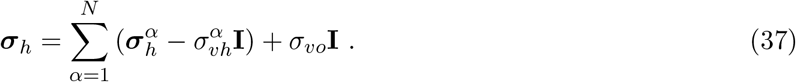

#### Remark 1.

For quasi-static incompressible hyperelastic behaviors, one generally satisfies mechanical equilibrium subject to boundary conditions and the incompressibility constraint *J* = 1. This set of equations determines the Lagrange multiplier *p* required to maintain incompressibility. In contrast, in the present G&R framework, one solves mechanical equilibrium (Eq. (7)) subject to boundary conditions and the mechanobiological equilibrium constraint **ϒ**_*h*_ = 1 (Eq. (6)). This set of equations determines the evolved homeostatic Lagrange multiplier *p_h_* consistent with a mechanobiologically equilibrated evolution. Note the difference between these two complementary *rate-independent* frameworks. The former satisfies the kinematic constraint *J* = 1, with *p* its associated stress-like Lagrange multiplier and *σ_υ_* to be determined. The latter satisfies the stress-like constraint **ϒ**_*h*_ = 1 (e.g., *σ_υ_* constrained via Eq. (35)), with *p_h_* an associated stress-like consistency parameter and *J_h_* to be determined. Thus, *p_h_* does not enforce incompressibility during mechanobiologically equilibrated evolutions; it yet enables a smooth transition to stresses associated with intermittent hyperelastic responses superimposed at a given G&R time *s = s_h_*, for which an evolved constraint *J = J_h_* is to be considered, see Appendix B.

#### Remark 2.

The pressure-sensitive *stimulus* function T in Eq. (33), which drives changes in volume via mass growth, resembles a pressure-sensitive *yield* function in rate-independent elastoplasticity, which triggers changes in volume via inelastic mechanisms but without mass exchange (such as for geomaterials, metallic foams, or filled polymers [22]). Since the present G&R formulation contains other noticeable differences when compared to classical elastoplasticity (e.g., mixture theory describing different behaviors of different constituents, not a homogenized material; pre-stresses with evolving natural configurations; and a multiplicative decomposition without explicit consideration of an elastic gradient), elastic predictor / inelastic corrector integration schemes typically used in computational plasticity [23, 24] do not seem well suited for integrating the present rate-independent constrained-mixture equations. Rather, we solved this set of nonlinear equations exactly using a different stress-point resolution procedure (Box 1), whose consistent linearization for efficient finite element implementations is addressed next. Of course, other resolution (or integration) schemes and consistent continuum (or algorithmic) linearizations may apply for other rate-independent G&R theories built upon different kinematic and / or constitutive assumptions (cf. [25]).

## 3. Consistent linearization of the continuum theory

In Section 2.5, we obtained (pre-)stresses for different constituents within a soft tissue in mechanobiological equilibrium by deriving, first, associated hyperelastic stresses during intermittent loading at a fixed G&R time *s = s_h_*, with *J = J_h_* and all constituent-specific natural configurations fixed, and, second, expressions specialized to a particular in vivo state **F** = **F**_*h*_ along the hyperelastic response path. During an incremental mechanobiologically equilibrated evolution, the soft tissue and ξ-constituent-specific natural configurations, with associated equilibrated stresses, evolve following different paths over (longer) G&R time scales [10]. We derive here tangent tensors for all load-bearing constituents and the mixture, consistent with a quasi-static G&R evolution of this type. Subsequent consideration of an evolving homeostatic Lagrange multiplier *p_h_*, arising from the mechanobiologically equilibrated constraint **ϒ**_*h*_ = 1, completes the (exact) consistent linearization of the formulation. Importantly, since numerical approximations (arising, typically, from integration of evolution equations) are absent in this formulation, the continuum and algorithmic linearizations coincide; a similar situation arises in computational hyperelasticity but not computational plasticity, where linearization of the resulting discrete equations is essential to preserve asymptotic rates of quadratic convergence characteristic of Newton’s methods [26].

### 3.1. Constituents that do not turnover

Equilibrated second Piola–Kirchhoff stresses in Eq. (26) can be written

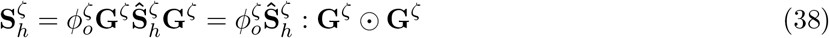

where the symbol : implies the usual contraction of two indices and 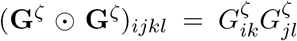, with **G**^*ζ*^ symmetric. Since **G**^*ζ*^ is constant, but 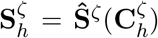 depends on **C**_*h*_ through 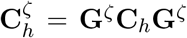, the constitutive (hyperelastic) tangent at the mixture level reads

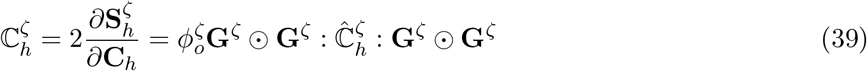

with 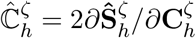 the associated tangent at the constituent level.

### 3.2. Constituents that turnover

Equilibrated second Piola–Kirchhoff stresses in Eq. (28) can be written, using Eq. (30),

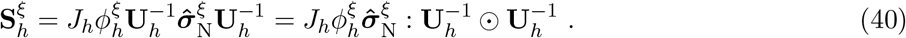

Since 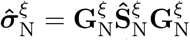 remains constant during mechanobiologically equilibrated evolutions, with natural configurations 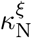 evolving accordingly, the (consistent) constitutive tangent at the mixture level reads

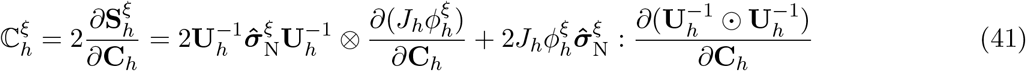

with ⊗ the usual dyadic product and the fourth-order tensor 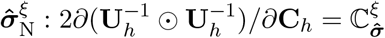, resulting from the double contraction between a second-order tensor 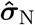 with a sixth-order tensor 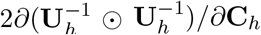, given in spectral decomposition form in Appendix D. Recalling from Section 2.2.2 that 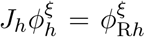 depends exclusively on *J_h_*, the chain rule on 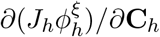, with 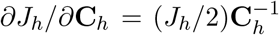, leads to

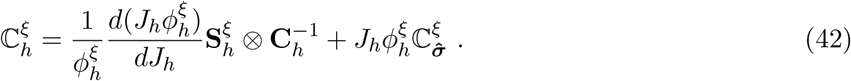

Note that Eqs. (11) and (12), each differentiated with respect to *J_h_*, constitute a linear system of *N^ξ^* equations with *N^ξ^* unknowns 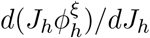, which can be solved explicitly. For example, if *η_ij_* = 1 ∀{*ξ_i_, ξ_j_*}, differentiation of the explicit result in Eq. (14) yields

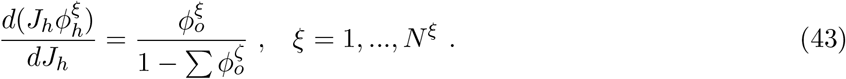

### 3.3. Lagrange multiplier contribution

The term 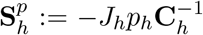 in Eq. (31) evolves consistent with the requirement in Eq. (33), that is 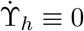, which defines *p_h_*(**C**_*h*_) constitutively during mechanobiological equilibrium and enables computation of a consistent tangent tensor

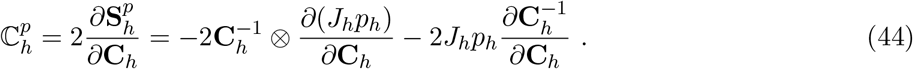

### 3.4. Mixture

Using Eq. (31), the referential tangent tensor for the mixture is

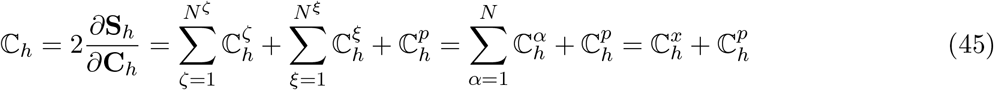

where all contributions are given above (Eqs. (39), (42), (44)). Its associated spatial tangent is given by the push-forward operation, from *κ_o_* to *κ_h_*,

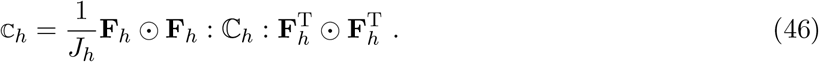

Linearization of the material model is summarized in Box 2.

#### Box 2: Mechanobiologically equilibrated constrained mixture model for G&R: Consistent linearization

i. Derivative of (referential) mass fractions for constituents *ξ* with respect to *J_h_* 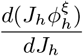, *ξ* = 1, …, *N^ξ^* determined explicitly from derivatives of Eqs. (11) and (12)
ii. Constitutive tangent for constituents *ζ* (mixture-level), which do not turnover

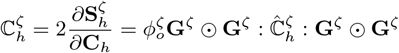

with constituent-level tangent 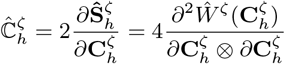
iii. Constitutive tangent for constituents *ξ* (mixture-level), which turnover

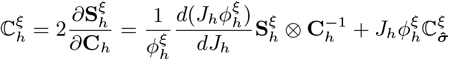

with 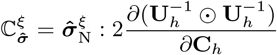 given in spectral form in Appendix D
iv. Contribution from evolving *p_h_* enforcing mechanobiologically equilibrated evolution

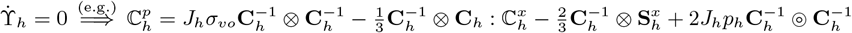
v. Referential and spatial tangent tensors for the mixture

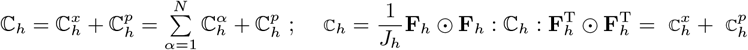

#### Example. (Cont.)

Equation (36), expressed in terms of mixture-level Lagrangian deformations and stresses, yields 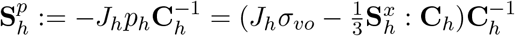 in Eq. (31). From Eq. (44)

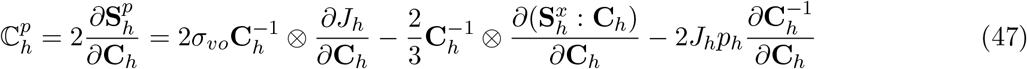

which, with 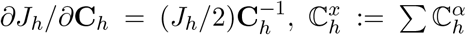 from Section 3.1 and 3.2, and 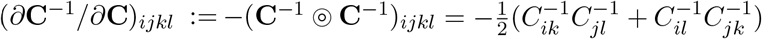, finally yields

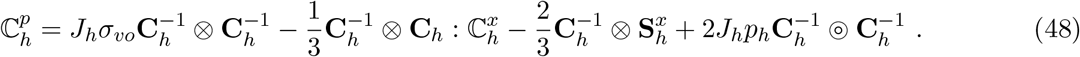

#### Remark 3.

The referential tangent tensor in Eq. (45), with Cartesian components 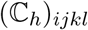, includes different contributions that lack the major symmetry *ijkl ↔ klij*, in general. First, the tensor 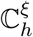 in Eq. (42) was derived from mechanobiological equilibrium considerations at the constituent level, that is 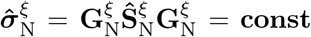 in Eq. (40), so its symmetry is not guaranteed. This is in contrast with the tensor 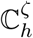 in Eq. (39), which derives from a hyperelastic potential, and hence is symmetric, even during G&R evolutions. Second, linearization of the consistency parameter *p_h_* in 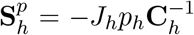 may yield additional contributions lacking major symmetry, for example, the second and third terms in the right-hand side of Eq. (48), because the extra stresses 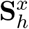 in Eq. (36) are not purely deviatoric, in general. This is in contrast with the first and fourth terms in Eq. (48), which are symmetric. All these non-symmetric tensors (cf. previous expressions and Appendix D as well) preserve the minor symmetries *ijkl ↔ jikl ↔ jilk ↔ ijlk*, which allows one to store their components for computational analyses using 6 × 6 matrices. This was not an issue for illustrative computations below, which required a non-symmetric finite element formulation consistent with the application of follower forces (pressure) that change direction as a body deforms (cf., for example, [27, 28]). Other constitutive formulations based on internal variables, such as finite kinematic growth [21], experimentally motivated non-associative plasticity [22], or fully (elastic and viscous) anisotropic finite strain nonlinear viscoelasticity [29], may similarly lead to non-symmetric consistent tangents for finite element implementations.

## 4. Original (loaded, in vivo) homeostatic state

The formulation outlined in Boxes 1 and 2 is intended to yield quasi-static G&R evolutions of threedimensional solids under continuous states of mechanobiological equilibrium. The formulation is defined locally at arbitrary material points and, hence, can be used to solve general boundary value problems employing standard finite element procedures. Inasmuch as **ϒ**_*h*_ depends on changes in stress relative to baseline values, one needs to pre-compute the original in vivo state, that is, an initially loaded state under mechanical and mechanobiological equilibrium. If the soft tissue behaves elastically under transient loads, then we can compute this initial state, subject to initial loads, by departing from a given initial geometry, close to the one at G&R time *s* = 0, and solving a boundary value problem with a nearly incompressible hyperelastic response. The (pre)stress computed at each material point after this initial, required calculation serves as a reference for the subsequent G&R computations by means of the specified equilibrium condition in **ϒ**_*h*_ = 1, as, for example, the local preservation of volumetric Cauchy stresses at the mixture level specified in Eq. (35).

Hence, consider the stresses derived in Appendix B and particularized to the initial in vivo state *κ_o_*, see Eq. (74), with *p* an actual Lagrange multiplier associated with the incompressibility constraint *J* =det **F** = 1 (i.e., at G&R time *s* = 0). In finite element implementations, however, one usually imposes near incompressibility through a penalty volumetric strain energy function *U*(*J*) (with an associated bulk modulus far exceeding the shear modulus), from which −*p*(*J*) is approximated by *dU*(*J*)/*dJ*, with *J* ≃ 1. For convenience, we consider a penalty function *U*(ln *J*) and approximate −*J_p_*(*J*) in Eq. (74) as *dU*(ln *J*)/*d*ln *J* =: *U*′(ln *J*), whereupon

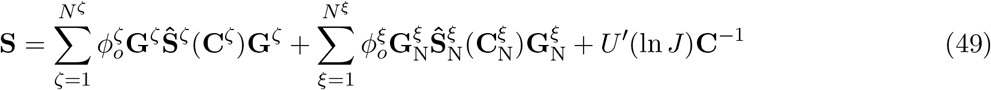

where 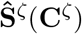 and 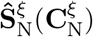 are computed from respective constituent-specific strain energy functions, see Eq. (72), with 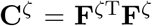 and 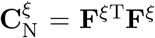 along with **F**^*ζ*^ = **FG**^*ζ*^ and 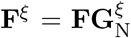. This expression for stresses is consistent with a constrained mixture theory for hyperelastic soft tissues with reference configuration *κ_o_* [30]. The constitutive tangent 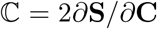 during this initial, elastic pre-loading step reads

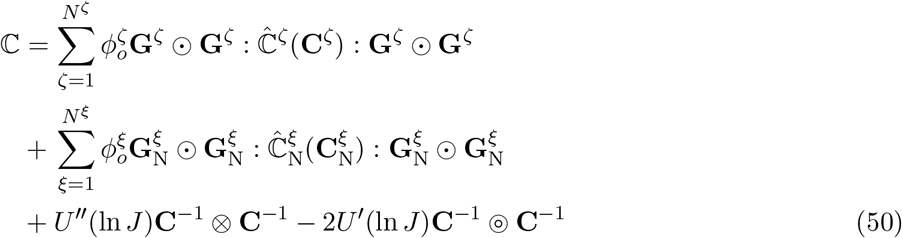

which can be implemented readily in a material subroutine for finite element analyses.

A step-by-step algorithm, including computations of associated stress and tangent moduli during both stages of our formulation (i.e., an initial pre-stressed state and subsequent equilibrated G&R evolution), is summarized in Box 3.

### Box 3: Two-stage algorithm to compute mechanobiologically equilibrated G&R evolutions

#### Stage I: Hyperelastic pre-loading to the original homeostatic state

Given (properties defined pointwise, in general):

- Geometry, external loads, and boundary conditions at G&R time *s* = 0
- Mass fractions 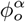 for all constituents *α* at G&R time *s* = 0
- Deposition stretch tensors **G**^*ζ*^ and 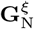 for all constituents *ζ* and *ξ* at G&R time *s* = 0
- Strain energy functions 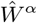 for all constituents *α*
- Volumetric (penalty) strain energy function *U*(ln *J*) for the mixture

##### (1.1)

Solve boundary value problem at *s* = 0, with:

Stresses and constitutive tangents in Eqs. (49) and (50)

##### (1.2)

Store at integration points (for the only time, for a smooth transition to Stage II):

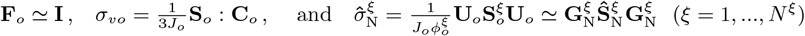

#### Stage II: Mechanobiologically equilibrated G&R evolution

Given, additionally (properties defined pointwise, in general):

- Change in external loads and/or other stimuli over G&R time *s* > 0
- *N^ξ^* − 1 independent gain-removal ratios *η^ij^* between constituents {*ξ_i_, ξ_j_*}

##### (II.1)

Solve boundary value problem at each *s = s_h_*, with:

Stresses and consistent tangents in Boxes 1 and 2

## 5. Illustrative numerical examples

Consider particular relations that have proven useful in describing vascular G&R, particularly for aneurysmal enlargement [12], cerebral vasospasm [31], aortic maladaptation in hypertension [15], and even the in vivo development of tissue engineered constructs [32]. A “four-fiber family” model includes three contributions: a stored energy function for an amorphous elastin-dominated matrix (*ζ ≡ e*)

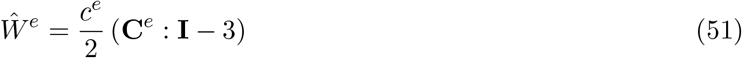

where *c^e^* is a shear modulus, plus circumferentially oriented passive smooth muscle (*ξ ≡ m*) and circumferentially, axially, and diagonally oriented (with angle α_0_ with respect to the axial direction) collagen fibers (*ξ ≡ c*) described by

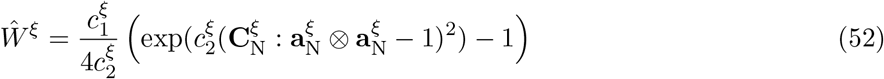

with 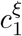 and 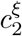 material parameters and 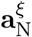 unit vectors along fiber orientations in 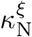. Effects of other constituents (such as proteoglycans) are captured phenomenologically via regression-based fits to data [15]. In addition, removal of constituents *ξ = m, c* is governed via a first-order type of kinetic decay, with *k^ξ^* rate parameters [7]. Finally, recall that net mass production is given by Eq. (4) via the removal and stimulus functions, here driven quasi-statically through Eq. (6). Table 1 lists all needed geometric and material parameters for an illustrative mouse aorta, noting that arteries tend to preserve their overall mass density *ρ* = 1050kg/ m^3^.

**Table 1:**
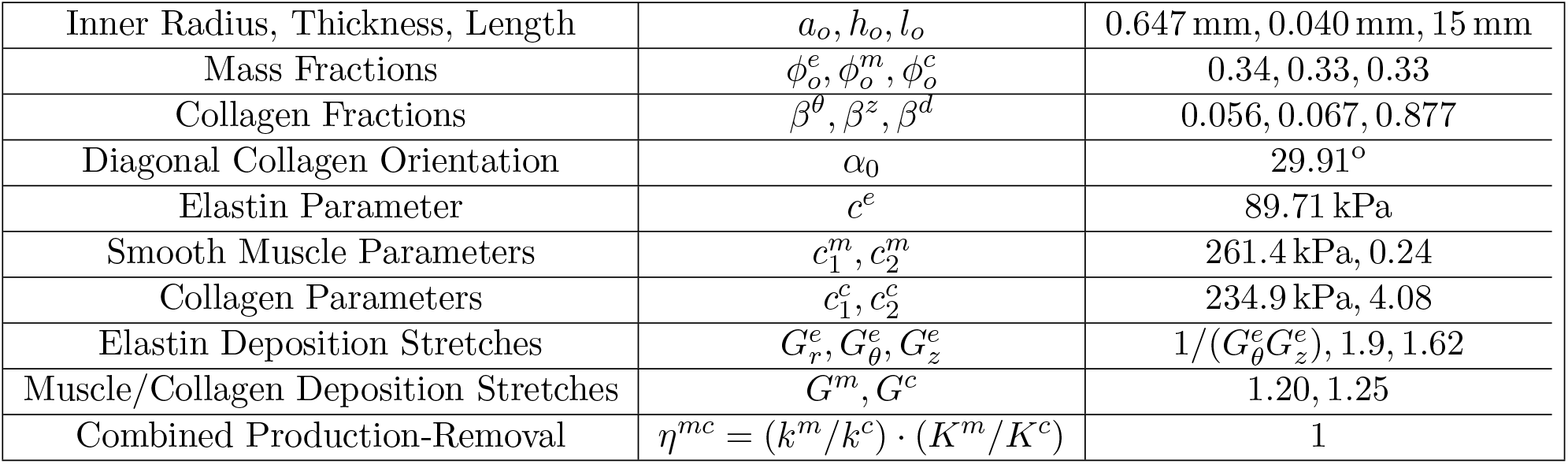
Representative baseline model parameters for a mouse descending thoracic aorta, assuming that elastin does not turnover during the G&R period but smooth muscle and collagen turnover continuously with constant deposition stretches (adapted from original homeostatic parameters in [15], with *η^mc^* = 1 because gain and rate parameters were determined while including inflammatory effects, which we do not consider here). Collagen family mass fractions are defined by 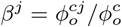, with *j* = *θ*, *z*, *d* representing circumferential, axial, and symmetric diagonal directions. The angle *α*_0_ is defined with respect to the axial direction. Additional values for 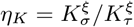 in Eq. (77) are given in the examples when appropriate.

|  |  |  |
| --- | --- | --- |
| Inner Radius, Thickness, Length | $a_o, h_o, l_o$ | 0.647 mm, 0.040 mm, 15 mm |
| Mass Fractions | $\phi_o^e, \phi_o^m, \phi_o^c$ | 0.34, 0.33, 0.33 |
| Collagen Fractions | $\beta^\theta, \beta^z, \beta^d$ | 0.056, 0.067, 0.877 |
| Diagonal Collagen Orientation | $\alpha_0$ | 29.91° |
| Elastin Parameter | $c^e$ | 89.71 kPa |
| Smooth Muscle Parameters | $c_1^m, c_2^m$ | 261.4 kPa, 0.24 |
| Collagen Parameters | $c_1^c, c_2^c$ | 234.9 kPa, 4.08 |
| Elastin Deposition Stretches | $G_r^e, G_\theta^e, G_z^e$ | $1/(G_\theta^e G_z^e), 1.9, 1.62$ |
| Muscle/Collagen Deposition Stretches | $G^m, G^c$ | 1.20, 1.25 |
| Combined Production-Removal | $\eta^{mc} = (k^m/k^c) \cdot (K^m/K^c)$ | 1 |

We implemented the present rate-independent constrained-mixture framework for G&R (Boxes 1 to 3) as a user material plugin in the open source software FEBio [33]. Because there is no need for local integration to update internal variables between incremental steps or storing long-term history-dependent variables, the implementation is straightforward and, from a numerical viewpoint, resembles that for a hyperelastic material (except for the non-symmetric consistent tangent and the need to solve numerically a system of equations for the Jacobian-dependent mass fractions *ϕ*^ξ^(*J*)). Solutions proceed in two stages. Stage I is not particular to the present formulation for G&R, but to any formulation that requires a prestressed initial configuration, such as hyperelastic [34] or G&R [35] formulations. Also note that mixture volumetric and constituent stresses are only stored once, at the end of Stage I, hence these initially computed variables play the role of (local) material parameters for Stage II. Importantly, however, the current version of FEBio used here (2.8.2) did not support materials with tangents having only minor symmetries, hence we added a new class of matrices and associated algebraic operators. In addition, some procedures within the FEBio source code were extended to handle non-symmetric constitutive tangents (i.e., an output from our material plugin) and FEBio was re-compiled accordingly.

### 5.1. Computation of the original homeostatic state (Stage I)

We compute here an original in vivo homeostatic state that will serve as the reference for all subsequent numerical simulations. Hence, consider an initially cylindrical arterial segment with inner radius *a_o_*, thickness *h_o_*, and length *l_o_* (Table 1) as a 3D finite element (FE) geometry (shown partially in Figure 2(a)). We fix axial displacements at both ends, with rigid body motions suppressed. As an external (pre-)load, we apply an in vivo value of blood pressure on the inner surface of the cylinder. Using a simple analytical (uniform) solution, see Appendix E or [7], one finds an in vivo pressure *P_o_* = 13.98 kPa = 104.9 mmHg consistent with model parameters in Table 1, as well as an original homeostatic Lagrange multiplier *p_o_* = 10.21 kPa. Hence, consistent with the concept of prestretch (and associated prestress), we define a volumetric function 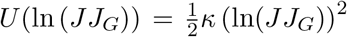, with *κ* = 10^3^*c^e^* a (penalty) bulk modulus and *J_G_* a pre-Jacobian satisfying *U*′(ln *J_G_*) = *κ* ln *J_G_* = −*p_o_*, that is, *J_G_* = exp(−*p_o_/κ*) for *J* = 1. With this, local stresses computed by Eq. (49) during the pre-loading stage are numerically close to those given analytically, the main difference being that the FE solution is appropriately non-uniform through the wall thickness. Only one static load step (with two-to-three global Newton–Raphson iterations) was needed to find associated equilibrium solutions (to machine precision) for the preload considered using different FE meshes. In any case, the computed equilibrium displacements for Stage I in Box 3 were negligible with respect to any characteristic length of the geometrical model, with **F** = **F**_*o*_ ≃ **I** and *J = J_o_* ≃ 1 as desired; compare initial (*a*, input) and computed (*b*, output) cross-sections in Figure 2. Furthermore, note that we do not prescribe an in vivo axial stretch 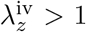 on this geometry (as one would do experimentally given an excised, unloaded vessel) because the computed solution implicitly includes an in vivo axial force (born by the axial constraints) consistent with the deposition stretches and associated stresses, in Table 1, which in turn allows us to refer deformations to this loaded configuration (i.e., λ_*z*_ = λ_*zo*_ = 1 refers to the original homeostatic state).

**Figure 2:**
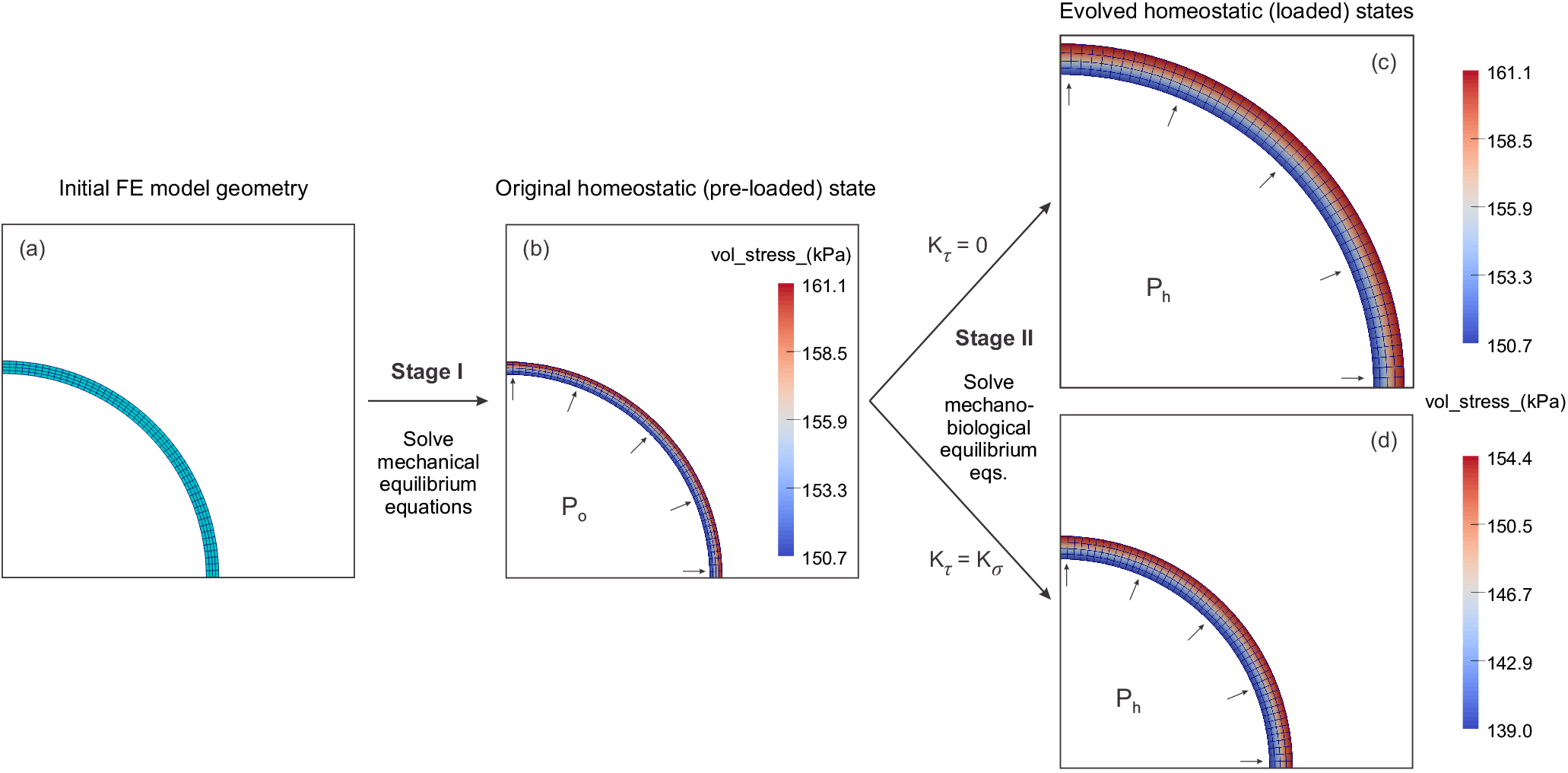
Initial (a) and computed after Stage I (b) quarter cross-section geometries (and meshes) for the initially cylindrical arterial segment considered in all FE simulations. Shown in (b), too, is the volumetric stress contour plot for this triaxially pre-stressed state consistent with parameters in Table 1, inner pressure *P_o_* = 13.98 kPa, flow rate ratio *ϵ_o_* = 1, and (implicit, with respect to an unloaded state) in vivo axial stretch 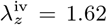. In addition, deformed cross-sections and volumetric stress contour plots are shown for increases in pressure up to *P_h_* = 1.6*P_o_* without (panel c, *K_xτ_*/*K_σ_* = 0) or with (panel d, *K_τ_*/*K_σ_* = 1) flow-induced shear stress effects on G&R (see Appendix C). Note the excessive dilation and thickening for the simulation (c) that preserves local values of volumetric stress with respect to (b) but neglects shear stress effects, see Eq. (35); in contrast, observe the mechano-adaptive thickening with little dilation (d), with slightly decreased volumetric stresses consistent with shear stresses reduced with respect to (b), see Eq. (79). Shown are smoothed stresses computed from (constant, but monotonic through the thickness) elemental stresses.

**Figure 3:**
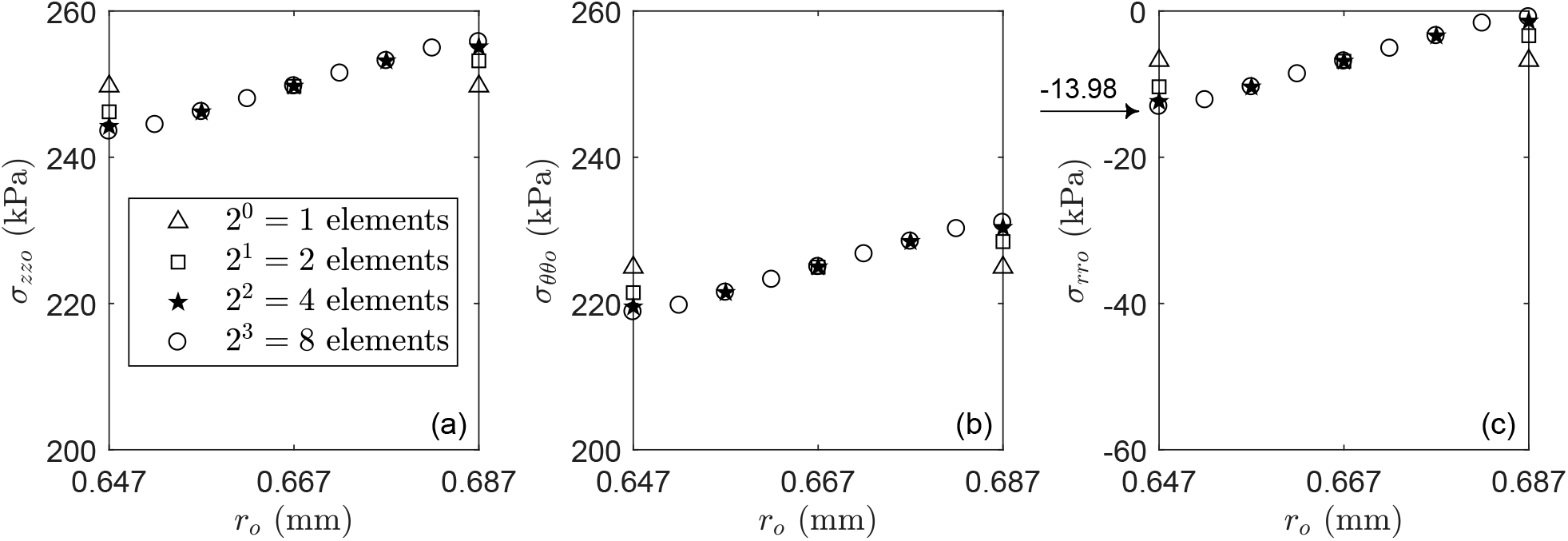
Mesh convergence study during the Stage I computation of the original pre-stressed homeostatic configuration as a function of the number of elements through the thickness of a mouse DTA (*a_o_* ≤ *r_o_* ≤ *h_o_,* Table 1) at a homeostatic state (*P_o_* = 13.98kPa, denoted by an arrow in panel c, with 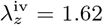). Four transmural elements were sufficient for this uni-layered wall model and thus were used in all subsequent simulations. Shown are axial (a), circumferential (b), and radial (c) Cauchy stresses at nodes interpolated from (constant, but monotonic through the thickness) elemental stresses. Note the different scales on the ordinates for the 40-micron thick wall.

We next conducted a mesh convergence study to determine an appropriate description of in vivo through-the-thickness stresses (3). After using different types of finite elements and discretizations, and because of the presence of pre-stresses (that tend to homogenize the stress field across the wall) and a relatively low thickness-to-radius ratio in the mouse aorta, we found that four displacement-based hexahedral elements give an accurate through-the-thickness solution for the present unilayered model. Figure 2(b) shows associated in vivo volumetric Cauchy pre-stresses *σ_υo_* = (1/3) tr***σ_o_***. This discretization is consistent with the one for displacements adopted in [36], where twelve mixed elements, linear in displacements and constant in pressure, were used to discretize three layers through the wall. As we explain below, volumetric locking is not expected during our quasi-static computations of mechanobiologically equilibrated G&R during Stage II, hence, mixed formulations with additional interpolations of pressure at the element level are not needed.

After computing this equilibrium solution, we stored total volumetric stresses for the mixture *σ_υo_* = (1/3*J_o_*)**S**_*o*_ : **C**_*o*_ and rotated Cauchy stresses for smooth muscle and collagen families 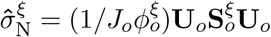 at integration points, which were needed (as reference variables) subsequently to compute mechanobiologically equilibrated G&R evolutions (Box 3). Note that we must compute these local values consistent with the non-uniform FE solution obtained, hence incorporating the extent of **F**_*o*_ − **I**, rather than from exact theoretical expressions (e.g., Eq. (30)) consistent with the exact reference limit **F**_*o*_ = **I**. Thus, in practice, one needs to store **F**_*o*_ and then compute G&R deformations during Stage II via 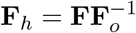 and 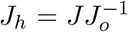. This ensures a smooth numerical transition between Stages I and II, that is, when **F** = **F**_*o*_ (i.e., 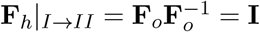 and 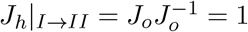, whereupon G&R starts right when Stage II does).

### 5.2. Equilibrated G&R response (Stage II) to increase in pressure, flow rate, or axial stretch

We verify here that mechanobiologically equilibrated responses computed for cylindrical geometries (that remain cylindrical) with the proposed FE formulation agree with responses computed with our previous theoretical formulation for thin-walled cylindrical arteries [7]. We note that the analytical rate-independent formulation was already verified and validated against a full integral heredity-based constrained model in [7] for long-term equilibrium solutions, and in [10] for slow loading relative to G&R.

For uniform material properties, the computed solutions are uniform in the circumferential and axial (in-plane) directions, but non-uniform along the radial direction consistent with different traction (pressure) conditions on the inner and outer boundaries. As explained in Appendix C, we can include a uniform flow-induced shear stress stimulus for mass production in the equilibrated function **ϒ**_*h*_ in Eq. (33); that is, we can include mechanobiological effects of a shear stress that is negligible mechanically (five orders of magnitude smaller [18]). Consideration of this augmented function will let us compute, first, pressure- and/or axial stretch-induced G&R responses for different intramural-to-shear stress gain ratios for mass production, and, second, flow-induced G&R responses. It will also let us compare FE outcomes against their corresponding analytical solutions, see Appendix C and Appendix E. Consistent with an axially uniform solution, we employ a single element along the axial direction and report deformations and band plots for a representative cross section.

Figure 2 also shows deformed cross-sections and contour plots of volumetric stress obtained for *P_h_* = 1.6*P_o_* = 22.38 kPa for two ratios of gain parameter, *K_τ_/K_σ_* = 0 (i.e., vanishing shear stress stimulus for mass production; mesh c) and *K_τ_/K_σ_* = 1 (i.e., both intramural and shear stress stimuli for mass production; mesh *d*). Figure 4 additionally shows associated pressure-dependent evolutions for global geometric parameters from the FE simulation (symbols) such as normalized inner radius (panel *a*) and thickness (*b*), as well as local variables at the mid-thickness such as volume ratio (*c*), and circumferential (*d*), axial (*e*), and volumetric (*f*) stresses, all compared to results computed with the uniform semi-analytical cylindrical model [7] (equivalently, Appendix E), which show excellent agreement. Note, however, the remarkably different responses that the mechanobiologically equilibrated models predict for *K_τ_/K_σ_* = 0 versus 1, which may be understood by examining the associated stimulus function **ϒ**_*h*_.

**Figure 4:**
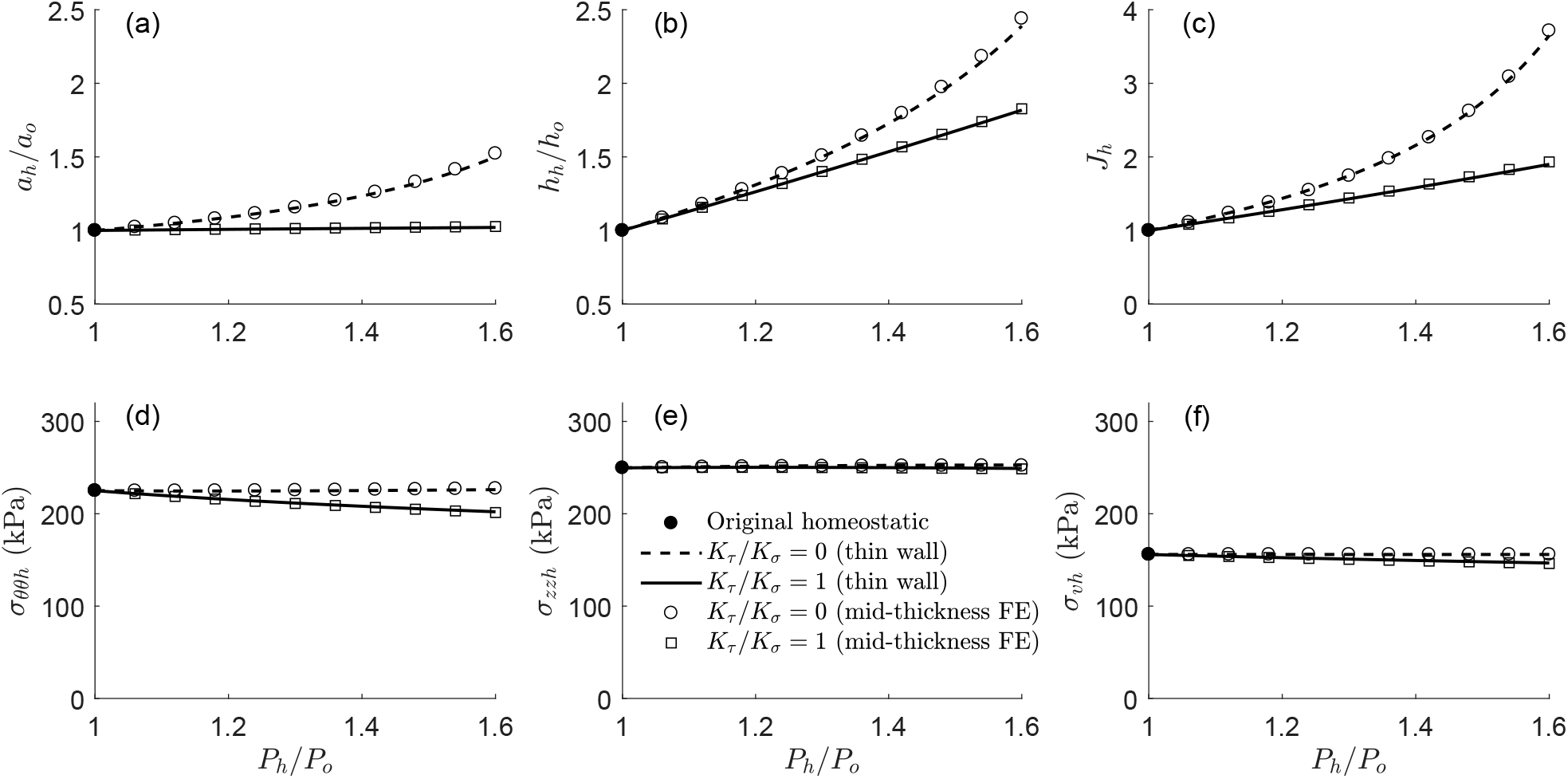
Analytical (thin-wall; lines) versus finite element (cylindrical model at mid-thickness; symbols) pressure-driven quasi-static G&R calculations for increases in pressure from *P_h_* = *P_o_* to *P_h_* = 1.6*P_o_* without (dashed line and open circles, *K_τ_*/*K_σ_* = 0) or with (solid line and open squares, *K_τ_*/*K_σ_* = 1) shear stress effects on G&R (see associated meshes and contour plots for *P_h_*/*P_o_* = 1.6 in Figure 2).

For *K_τ_/K_σ_* = 0, the constraint **ϒ**_*h*_ = 1 implies preservation of volumetric stresses *σ_υh_ = σ_υo_* (recall Eqs. (35) or (37)), which the FE simulation satisfies locally at each radial location (Fig. 2) and, in particular, at the mid-thickness (Fig. 4). Albeit shown at only the mid-thickness, circumferential and axial Cauchy stresses remain nearly constant during this quasi-static G&R evolution. Yet, this restoration of stress via mass growth arises from increases in inner radius (circumferential stretch) and thickness (radial stretch), and thus associated volume (Jacobian), which depart from the theoretical mechano-adaptive limits 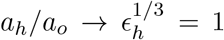 and 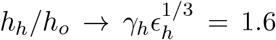 (when *ϵ_h_ = Q_h_/Q_o_* = 1 and *γ_h_ = P_h_/P_o_* = 1.6, with *Q* blood flow rate), cf. [18]. Indeed, the analytical model predicts a marked growth response (*a_h_/a_o_* → ∞ and *h_h_/h_o_* → ∞) for *K_τ_/K_σ_* = 0 and *P_h_* → 1.95*P_o_*, which suggests a static mechanobiological instability (or unbounded critical point) against increases in pressure, cf. [8]. Importantly, when accounting for the radially non-uniform distributions of stress / stretch, this instability developed earlier at the outer than at the inner layer of elements, which caused the FE simulation to diverge before reaching the theoretical blow-up pressure 1.95*P_o_* predicted by the uniform formulation. This finding suggests that full 3D simulations can be needed even for thin-walled arteries. It also reminds us of the need to account for all mechanobiological stimuli (endothelial cell shear and interstitial cell normal stresses) even when one component of stress is negligible mechanically.

In particular, for *K_τ_/K_σ_* = 1, the constraint **ϒ**_*h*_ = 1 introduces an additional restriction on changes in flow-induced wall shear stress *τ_υh_*, which must offset changes in volumetric stress *σ_υh_* (recall Eq. (79)). For example, for the idealized wall shear stress expression *τ_w_* = 4*μQ/πa*^3^ (with *μ* the blood viscosity) that we employ, a restriction on *τ_wh_* with preserved cardiac output controls luminal dilation, which in turn affects the equilibrium solution. The FE simulation then predicts a slight decrease in volumetric stress *σ_υh_* at each radial coordinate for *P_h_/P_o_* = 1.6 (Fig. 2), shown at the mid-thickness for different pressures in Fig. 4. For *K_τ_/K_σ_* = 1 and *P_h_/P_o_* = 1.6, *σ_υh_/σ_υo_ = τ_wh_/τ_wo_* ≈ 0.94 leads to an increase in inner radius *a_h_/a_o_* = (*τ_wo_/τ_wh_*)^1/3^ ≈ 1.02, much lower than for *K_τ_/K_σ_* = 0, and consistent with analytical solutions. The nearly preserved inner radius (circumferential stretch) and increase in thickness (radial stretch), with an associated increase in volume (Jacobian), closely follow the mechano-adaptive limits 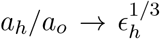 and 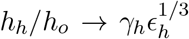. Yet, a slight, non-ideal over-thickening (~ 10%, due to the presence of elastin, which does not turnover and prevents perfect adaptations) was consistent with the slight decrease in due to a decrease in circumferential (~ 10%) rather than axial (which remained nearly constant) stress. Importantly, the pressure-driven mechanobiologically equilibrated (analytical and FE) solutions followed this trend up to *P_h_/P_o_* = 2, hence suggesting a mechanobiologically stable (bounded) quasi-static evolution within physiological limits [8].

Similar quasi-uniform FE simulations were performed for prescribed changes in flow rate *Q_h_ > Q_o_* (Figure 5) and axial stretch λ_*h*_ > λ_*o*_ (Figure 6) with or without shear stress effects on G&R. Like the pressure-driven simulations addressed above, values computed at the mid-thickness agreed well with the analytical calculations. In particular, an artery does not respond to increases in flow if *K_τ_* = 0, whereas it mechanoadapts for *K_τ_ = K_σ_* with, unlike the previous pressure-driven cases, an associated increase in volumetric stress consistent with a computed increase in flow-induced wall shear stress *τ_wh_/τ_wo_ = ϵ_h_*/(*a_h_/a_o_*) ≈ 1.05 for *ϵ_h_* = 1.6 (Figure 5). Consideration of shear stress effects to achieve mechanoadaptation was even more critical for isolated changes in axial stretch (i.e., with *ϵ_h_ = γ_h_* = 1). Indeed, Figure 6 shows that the artery increased its length *l_h_/l_o_* = λ_*zh*_ > 1 while nearly preserving inner radius (λ_*θh*_ ≈ 1) and thickness (λ_*rh*_ ≈ 1) if *K_τ_ = K_σ_*, hence *J_h_* > 1 (i.e., by net mass production), indicative of a mechanical adaptation in terms of intramural volumetric and wall shear stresses. Note, however, that axial stress necessarily increases consistent with the axially stretched (elastic) elastin, with circumferential stress regulated and nearly maintained consistent with the slight change in inner radius and thickness with constant blood pressure. In contrast, inner radius (λ_*θh*_ < 1) and thickness (λ_*rh*_ < 1) decreased for *K_τ_* = 0, which even led to respective decrements in volume *J_h_* < 1 (i.e., by net mass removal) for *l_h_/l_o_* = λ_*zh*_ > 1.2, indicative of a mechanical maladaptation by flow shear stress dysregulation along with excessive increments and decrements of axial and circumferential stresses, respectively, even if the volumetric stress is preserved. Again, the elastically stretched elastin, and its associated increase in axial stress at the mixture level, plays a central role in this lack of adaptation by G&R.

**Figure 5:**
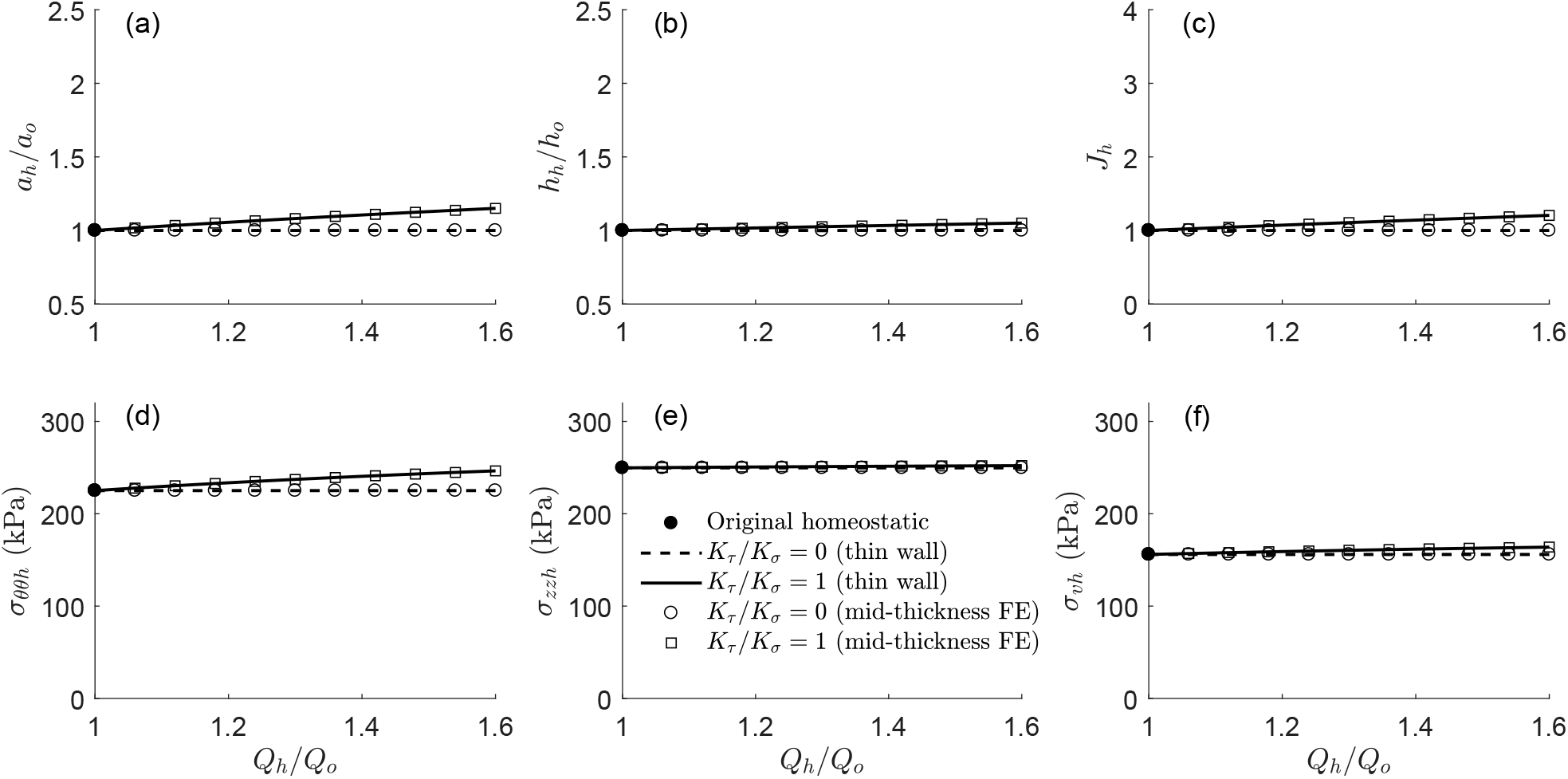
Analytical (thin-wall; lines) versus finite element (cylindrical model at mid-thickness; symbols) flow-driven quasistatic G&R calculations for increases in blood flow rate from *Q_h_* = *Q_o_* to *Q_h_* = 1.6*Q_o_* without (dashed line and open circles, *K_τ_*/*K_σ_* = 0) or with (solid line and open squares, *K_τ_*/*K_σ_* = 1) shear stress effects on G&R.

Although it was possible to compute these quasi-uniform solutions in single load steps during Stage II in Box 3, we nevertheless re-computed them with 10 incremental load steps (plus an initial one for Stage I; symbols in Figs. 4 to 6) to compare quasi-static evolutions with the analytical formulation. Table 2 shows associated quadratic rates of convergence for residual force and energy during global Newton–Raphson iterations, without line searches, and total simulation times (on a single CPU processor Intel^®^ Xeon^®^ E5 at 3 GHz in a Workstation Dell Precision 5810 with 32GB RAM). Note that the computational efficiency of this rate-independent G&R formulation is comparable to that obtained for hyperelastic computations, for which we used the same mesh, boundary conditions, applied loads, and incremental steps along with Eqs. (49) and (50), and, hence, is expected to be comparable as well to that obtained by efficient rate-dependent G&R formulations (e.g., in 2D FE analyses [37]). Lastly, because mechanobiologically equilibrated G&R is not constrained isochorically (recall Remark 1), no special formulations were needed to prevent volumetric locking during these computations, a numerical issue characteristic of nearly incompressible elastic or distortional elastoplastic responses [24, 38], but also transient elastic responses during G&R [36, 39]. In this respect, all solution variables changed gradually between adjacent elements and agreed well with analytical results, including global geometric outputs (e.g., inner radius and thickness).

**Table 2:** Asymptotically quadratic rates of convergence for residual force (left sub-columns) and energy (right sub-columns) during a typical global time step for the two axially uniform G&R simulations shown in Fig. 2 (c and d) as well as total elapsed CPU time (1 +10 time steps each, Fig. 4) using a single processor. Note that both convergence rate and computational time are comparable to those for a hyperelastic computation using the same FE model and increase in pressure without G&R (not shown).

| | G&R ($K_\tau/K_\sigma = 0$) | | G&R($K_\tau/K_\sigma = 1$) | | Hyperelastic | |
| --- | --- | --- | --- | --- | --- | --- |
|  | force | energy | force | energy | force | energy |
| Iteration 0 | 2.9E+00 | 1.5E-00 | 2.6E+00 | 9.0E-02 | 6.5E+00 | 5.1E-01 |
| Iteration 1 | 2.2E-01 | 9.7E-03 | 3.3E-02 | 5.1E-04 | 7.3E+01 | 6.8E-02 |
| Iteration 2 | 1.5E-04 | 1.4E-08 | 1.2E-05 | 4.3E-09 | 4.9E-01 | 2.3E-04 |
| Iteration 3 | 6.1E-10 | 1.7E-18 | 3.2E-10 | 3.6E-20 | 9.4E-04 | 4.7E-09 |
| Iteration 4 |  |  |  |  | 1.1E-10 | 1.7E-18 |
| Time (1+10 steps) | 6.63 s |  | 6.76 s |  | 6.37 s |  |

**Figure 6:**
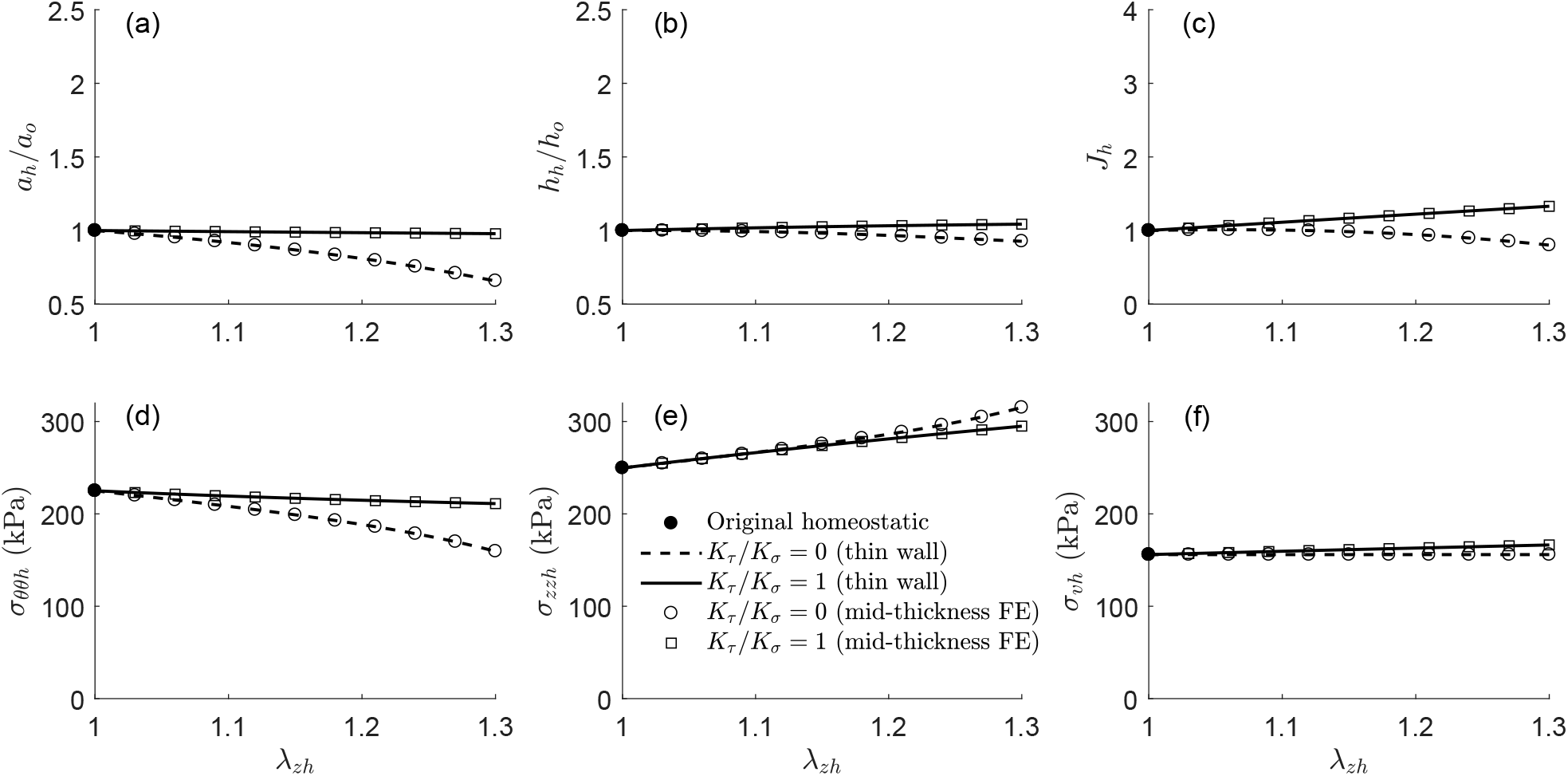
Analytical (thin-wall; lines) versus finite element (cylindrical model at mid-thickness; symbols) axial stretch-driven quasi-static G&R calculations for increases in axial stretch from λ_*h*_ = λ_*o*_ = 1 to λ_*h*_ = 1.3λ_*o*_ = 1.3 without (dashed line and open circles, *K_τ_*/*K_σ_* = 0) or with (solid line and open squares, *K_τ_*/*K_σ_* = 1) shear stress effects on G&R. Note the increase in axial stress contributed by the elastically stretched elastin.

### 5.3. Non-uniform equilibrated G&R response (Stage II) to localized degradation of elastin

Next, consider non-uniform deformations of an arterial segment. Since the idealized formula *τ_w_* = 4*μQ/πa*^3^ is only valid for fully developed steady flows in a long circular segment, we first considered G&R driven by intramural stresses only, for which *K_σ_* = 0 and *K_τ_* = 0, such that volumetric stresses were preserved locally, recall Eq. (35). As an example, a G&R response that can be modeled with *K_τ_* = 0 in **ϒ** is the enlargement of fusiform aneurysms, where the medial layer appears to be severely damaged (hence no shear stress regulation of smooth muscle tone) and enlargement likely occurs primarily via turnover of remnant collagen in the adventitia [12, 35, 40–43].

Since medial damage implies degradation of elastin laminae, consider the mechanobiologically equilibrated G&R analysis in Figure 7 performed for a uniform, progressive degradation of elastin, either with decreasing 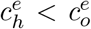 or an increasing damage parameter 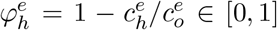, while preserving inner pressure *P_h_ = P_o_*, cf. [8]. Due to the close agreement between the uniform analytical and (quasi-)uniform FE simulations shown in Fig. 7, for both *K_τ_/K_σ_* = 0 and a slight *K_τ_/K_σ_* = 0.1, these results provide important guidance for the interpretation of more complex FE simulations. In this case, uniformly degraded elastin, without shear stress effects on G&R, can eventually lead to an unbounded G&R response, specifically for 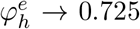 (Fig. 7, *K_τ_/K_σ_* = 0), which suggests that locally degraded elastin coupled with endothelial dysfunction could associate with local aneurysmal expansions that may develop quasistatically but remain bounded because constraints on lateral displacements prevent a local asymptotic enlargement. To test this hypothesis within our rate-independent G&R framework, consider the same initial cylindrical geometry (hence, the same homeostatic state after Stage I), although discretized with 60 elements along the axial direction to enable finer non-uniformities within the central region (see meshes (a) in Figure 8, computed after Stage I). Subsequently, we locally increase 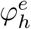 at selected regions to drive quasi-static G&R FE simulations. This way, we model effects of elastin degradation on fully resolved G&R, not the underlying cause of this degradation, for which one would need to couple a damage model with the present rate-independent formulation.

**Figure 7:**
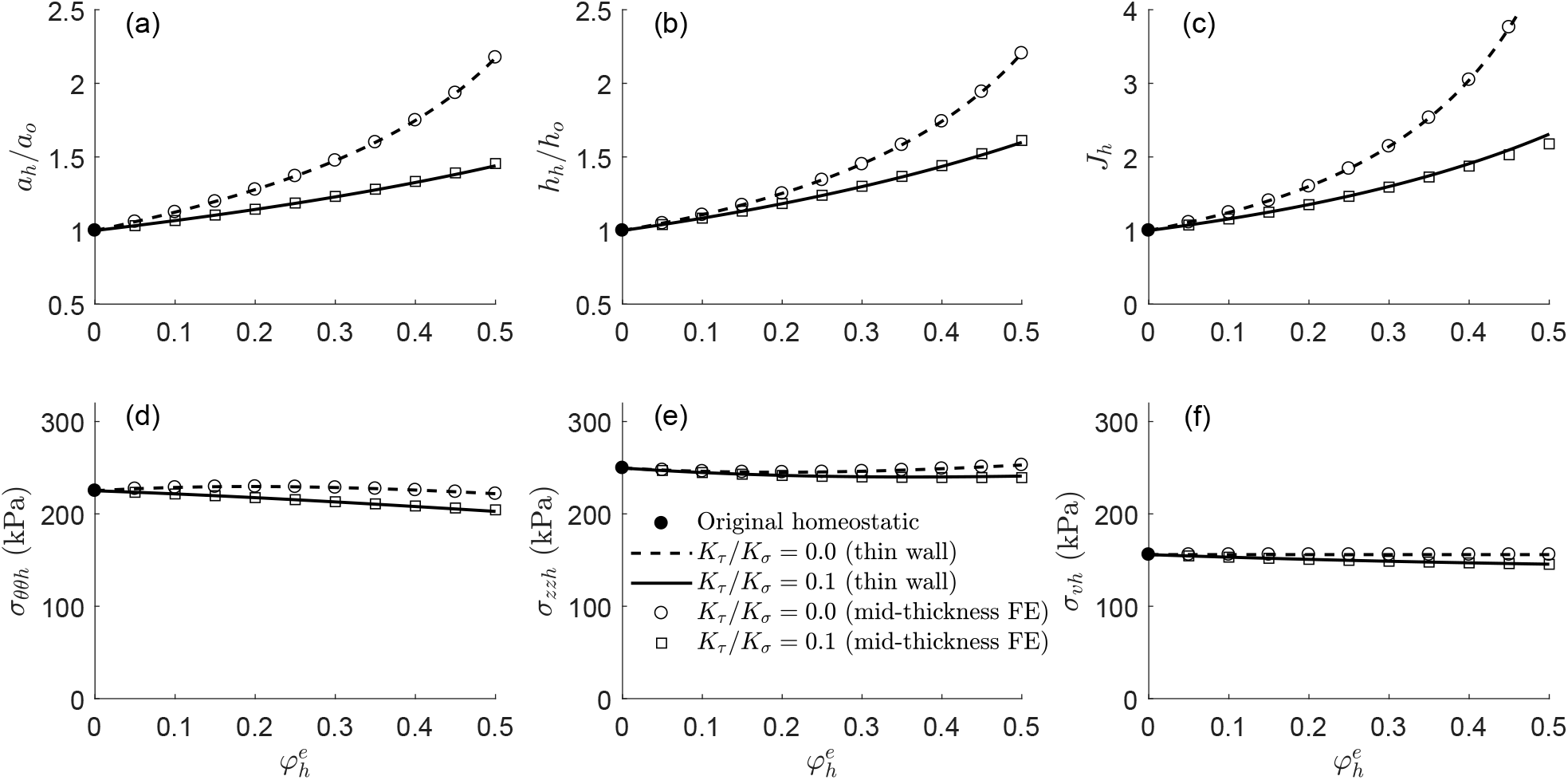
Analytical (thin-wall; lines) versus finite element (cylindrical model at mid-thickness; symbols) elastin degradation-driven quasi-static G&R calculations for increases in the degradation parameter 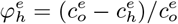 from 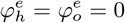 up to 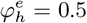, while preserving inner pressure *P_h_* = *P_o_*, without (dashed line and open circles, *K_τ_*/*K_σ_* = 0) or with slight (solid line and open squares, *K_τ_*/*K_σ_* = 0.1) shear stress effects on G&R. Both models predict asymptotic thickening and dilation with moderate change in biaxial stress for 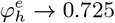 when *K_τ_*/*K_σ_* = 0 (not shown), cf. [8].

**Figure 8:**
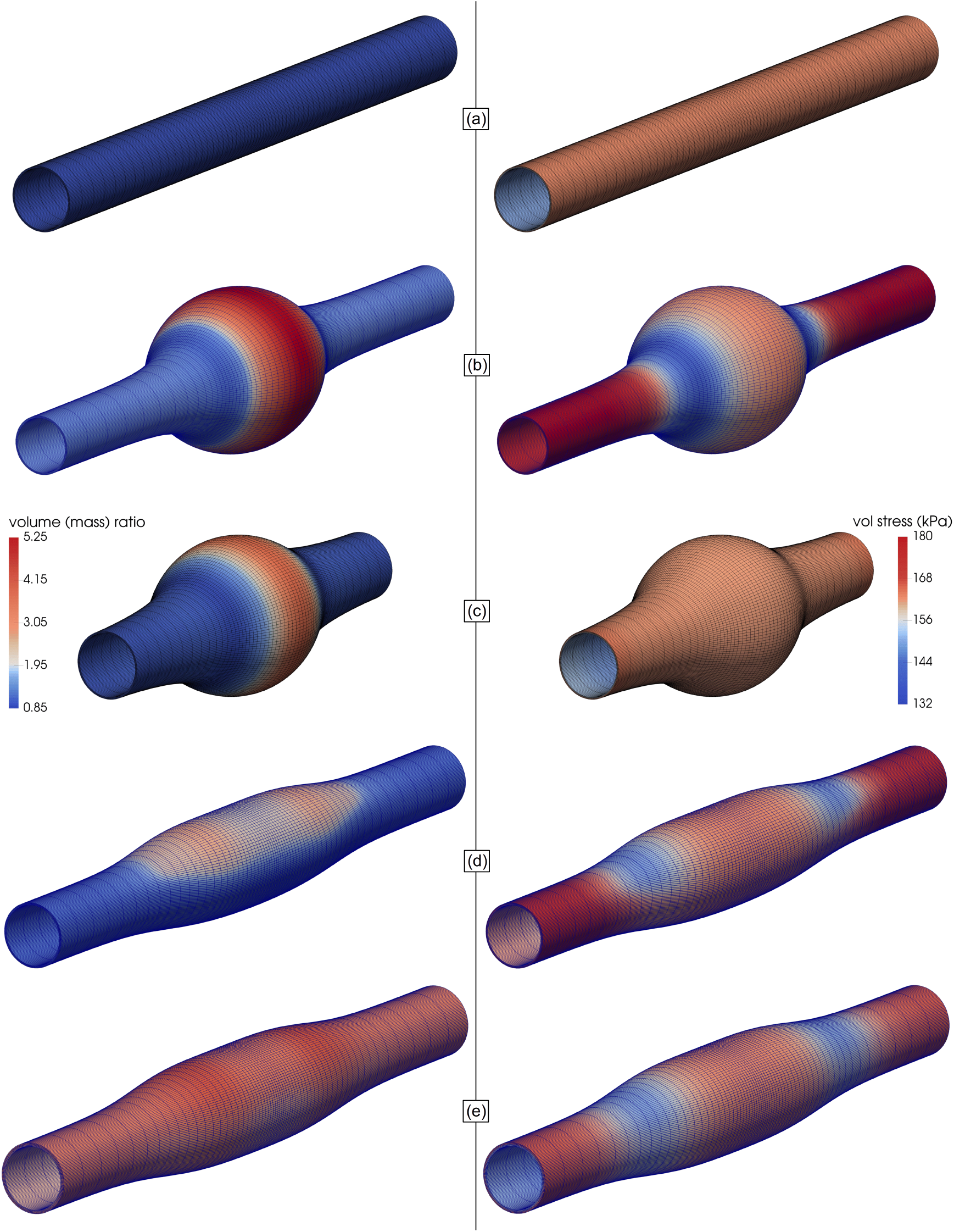
Initial (after Stage I; a) and computed (in Stage II; b to e) mechanobiologically equilibrated states for an arterial segment with axisymmetrically (b, c) or asymmetrically (d, e) prescribed elastin damage given either prescribed axial displacements (b, d, e) or force (c) at the ends, as well as normotensive (b to d) or hypertensive (e) conditions. Shown are deformed meshes and contour plots of the Jacobian deformation (left column) and volumetric Cauchy stress (right column) for: 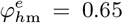, *z*_*o*d_ = *l_o_*/4, and *ν_z_* = 2 in Eq. (53), with *K_τ_*/*K_σ_*(*z*_*o*min_) = 0.35 in Eq. (54), for *P_h_* = *P_o_*; (c) 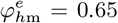, *z*_*o*d_ = *l_o_*/4, and *ν_z_* = 2 in Eq. (53), with *K_τ_*/*K_σ_* = 0, for *P_h_* = *P_o_*; (d) 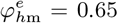, *z*_*o*d_ = *l_o_*/3, *θ*_*o*d_ = π/3, and *ν_z_* = *ν_θ_* = 5 in Eq. (55), with *K_τ_*/*K_σ_*(*z*_*o*min_) = 0.35 in Eq. (54), for *P_h_* = *P_o_*; and (e) 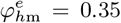 = 0.35, *z*_*o*d_ = *l_o_*/3, *θ*_*o*d_ = π/3, and *ν_z_* = *ν_θ_* = 5 in Eq. (55), with *K_τ_*/*K_σ_*(*z*_*o*min_) = 0.35 in Eq. (54), for *P_h_* = 1.5*P_o_*. The FE model comprised *N_r_* × *N_θ_* × *N_z_* =4 × 160 × 60 = 38400 displacement-based linear hexahedral elements.

With preserved inner pressure *P_h_ = P_o_*, consider two different spatial distributions for 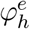 in terms of cylindrical coordinates *r_o_* ∈ [*a_o_, a_o_ + h_o_*], *θ_o_* ∈ [0, 2*π*], and *z_o_* ∈ [*z*_*o*min_, *z*_*o*max_] in the reference homeostatic configuration *κ_o_*, where *z*_*o*min_ and *z*_*o*max_ = *z*_*o*min_ + *l_o_* are minimum and maximum axial coordinates. Consider first an axisymmetric degradation localized at *z*_*o*m_ = (*z*_*o*max_ + *z*_*o*min_)/2 that diminishes gradually with |*z_o_* − *z*_*o*m_|, namely

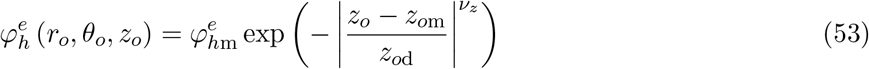

with *ν_z_* > 0 and *z*_*o*d_ > 0 respective axial exponential decay and deviation parameters. Moreover, 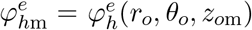 describes the maximum local degradation at a given load Step, and 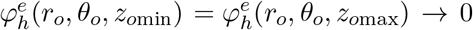 at both ends. Also, 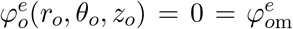 throughout the arterial wall at the original homeostatic state. Then, for constant blood pressure, the FE simulation can be advanced in a quasi-static (time-independent) manner by increasing a single scalar 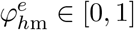 over subsequent steps.

In a first attempt to reproduce this local enlargement of the artery, we prescribed *K_τ_* = 0 throughout. Because axial displacements are fixed at the boundary ends, and a circumferential expansion with an overall axial retraction was observed at the central region of the artery, local axial stretching is expected next to the ends. Recalling the G&R response against increases in axial stretch with *K_τ_* = 0 (Figure 6), one would expect a constriction of the artery close to the ends of the model. Albeit not shown, this abnormal response was verified numerically with our FE model, which suggested the need for a shear stress effect on G&R, with associated inner radius regulation, in the non-damaged areas. Hence, as a first approximation (in the absence of a fluid-solid interaction solution), we simulated this effect by letting *K_τ_* > 0 (along with *τ_w_* = 4*μQ/πa*^3^, recall Eq. (79)) next to the lateral ends, which yet remain cylindrical, but *K_τ_* = 0 in a central damaged area where the aneurysm develops. To do so, we considered an axial distribution for *K_τ_/K_σ_* similar to that for elastin degradation in Eq. (53), namely

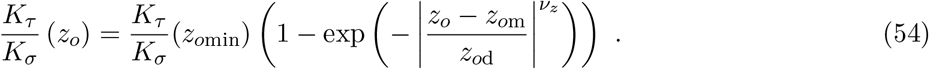

Thus, *K_τ_/K_σ_*(*z*_*o*min_) = *K_τ_/K_σ_*(*z*_*o*max_) > 0 is maximum at both ends and *K_τ_/K_σ_*(*z*_*o*m_) = 0 at the central cross-section. Meshes (b) in Figure 8 show the deformed configuration and associated contour plots of the volume ratio (left) and volumetric stress (right) for 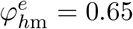, *z*_*o*d_ = *l_o_*/4, = 2, and *K_τ_/K_σ_*(*z*_*o*min_) = 0.35 in Eqs. (53) and (54), which forms an axisymmetric fusiform aneurysm. Importantly, while the uniform formulation, with 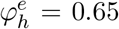 and *K_τ_* = 0, predicts a nearly isotropic expansion within a given cross-section, with λ_*θh*_ = *a_h_/a_o_* = 3.97 and λ_*rh*_ = *h_h_/h_o_* = 4.17, and λ_*zh*_ = *l_h_/l_o_* = 1 prescribed (cf. Figure 7), the non-uniform 3D FE simulation predicts a different expansion at the central cross-section *z*_*o*m_, with λ_*θh*_ = 2.50 ≃ *a_h_/a_o_* and λ_*rh*_ = 1.44 ≃ *h_h_/h_o_* at the mid-thickness, but λ_*zh*_ = 1.50 = *l_h_/l_o_*. Furthermore, lateral, radially unexpanded, cross-sections yet cause the aneurysm to grow axially at *z*_*o*m_ (λ_*zh*_ = 1.50 vs. 1 for the uniform case) and prevent it from growing asymptotically, reducing growth along circumferential (λ_*θh*_ = 2.50 vs. 3.97) but especially radial (λ_*rh*_ = 1.44 vs. 4.17) directions, that is, reduced dilation but especially thickening. In contrast, mean in-plane mechanobiologically equilibrated stresses at *z*_om_ (*σ_θθh_* = 215 kPa and *σ_zzh_* = 259 kPa) remain close to the uniform predictions (*σ_θθh_* = 214 kPa and *σ_zh_* = 260kPa), cf. Figure 7. Note, too, that a marked axial retraction predicted at the shoulders of the aneurysm contributes to an overall retraction of the central region. Hence, as mentioned above, finite elements close to the boundaries *z*_*o*min_ and *z*_*o*max_ are highly stretched axially (reaching up to λ_*zh*_ = 1.69), which justify the distribution for *K_τ_/K_σ_* in Eq. (54) to prevent radial constriction, to some extent, by wall shear stress regulation (recall the analysis in Figure 6, conducted up to a more moderate λ_*zh*_ = 1.3) at the expense of an increased volumetric stress *σ_υh_* in those regions (mesh (b)-right in Figure 8) consistent with an increased shear stress for constant cardiac output and reduced lumen. The volumetric stress then rapidly decreases in the transition zone where the inner radius starts to increase, consistent with a lower shear stress combined with some remnant *K_τ_/K_σ_* > 0, to finally recover the original homeostatic distribution for *σ_υo_* within the aneurysmal region (compare to mesh (a)-right), consistent with a vanishing shear stress effect for *K_τ_/K_σ_* = 0.

Indeed, it is instructive to consider a different set of boundary conditions at both ends *z*_*o*min_ and *z*_*o*max_ to analyze different responses of the nearby elements. Hence, to alleviate the constraint on axial displacements as well as the associated axial stretching of elastin and consequent radial constriction of the artery, we repeated the previous simulation while prescribing a constant axial force at the ends, coupled with *K_τ_* = 0 throughout the arterial segment. Meshes (c) in Figure 8 show the formation of an axisymmetric fusiform aneurysm similar to the previous one (b), though with a pronounced axial recoiling (*l_h_* = 9.85 mm vs. *l_o_* = 15 mm) that effectively alleviates the local axial stretching (λ_*zh*_ − 1) near the boundaries. Consistent with the analysis in Figure 6, the inner radius at the boundaries is preserved in this case even without considering shear stress effects on G&R. Also consistent with *K_τ_* = 0, and unlike the simulation (b), the distribution of *σ_υh_* is preserved with respect to the original homeostatic preloaded state (a). Equilibrium values at the apex reach, in this case, λ_*θh*_ = 2.41, λ_*rh*_ = 1.33, λ_*zh*_ = 1.11, *σ_θθh_* = 221 kPa, and *σ_zzh_* = 253 kPa, measured at the mid-thickness. Arguably, neither axial displacements nor force are necessarily fixed in actual aneurysms; hence, although these simpler boundary conditions served to illustrate limiting situations and radically different implications, more realistic boundary conditions are needed to simulate aneurysmal enlargement and associated implications.

Consider now a degradation localized at *z*_*o*m_ and *θ*_*o*m_ = *π* that diminishes gradually with |*z_o_ − z*_*o*m_| and |*θ_o_ − θ*_*o*m_|, namely

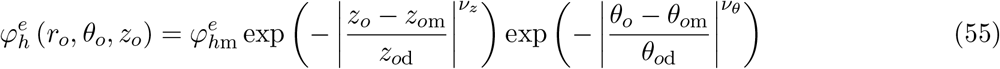

with *ν_θ_* > 0 and *θ*_*o*d_ > 0. Moreover, 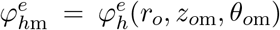 describes maximum local degradation at a given “load” step, 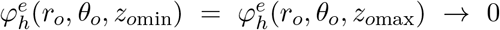 at both ends, and 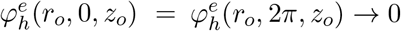. As before, we advance the quasi-static FE simulation, for constant inner pressure, by increasing a single scalar 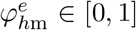 over subsequent steps. Meshes (d) in Figure 8 show the deformed configuration as well as contour plots for *J_h_* and *σ_υh_*, with 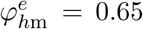, *z*_*o*d_ = *l_o_*/3, *θ*_*o*d_ = *π*/3, and *ν_z_ = ν_θ_* = 5 in Eq. (55). This yields an axially longer but radially asymmetric fusiform aneurysm. Axially non-uniform shear stress effects were included again, with *K_τ_/K_σ_*(*z*_*o*min_) = 0.35, *z*_*o*d_ = *l_o_*/3, and *ν_z_* = 5, in Eq. (54). Importantly, even if the maximum local value for 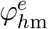 is the same as in simulations (b) and (c), the larger surrounding area of non-damaged material (in axial and circumferential directions) prevents the aneurysm from growing to a larger extent. In this case, both λ_*θh*_ = 2.29 > 1 and λ_*rh*_ = 1.35 > 1, but λ_*zh*_ = 0.65 < 1 at the apex (*z_o_ = z*_*o*m_, *θ_o_ = π*), where in-plane local stresses (*σ_θθh_* = 250 kPa and *σ_zzh_* = 224 kPa) yet remained bounded. Indirectly induced growth at the diametrically opposed location (*z_o_ = z*_*o*m_, *θ_o_* = 0) was lower, namely λ_*θh*_ = 1.17 > 1 and λ_*rh*_ = 1.30 > 1, but λ_*zh*_ = 0.78 < 1, where in-plane local stresses reached the same equilibrium values as at the apex.

Finally, let an asymmetric degradation of elastin as in Eq. (55), with the same distribution parameters *z*_*o*d_ = *l_o_*/3, *θ*_*o*d_ = *π*/3, and *ν_z_ = ν_θ_* = 5, as well as *K_τ_/K_σ_*(*z*_*o*min_) = 0.35 in Eq. (54) and simulation (d), develop in parallel with an increase in transmural pressure *P_h_* > *P_o_*. Meshes (e) in Figure 8 show deformed configurations and contour maps for 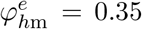 and *P_h_* = 1.5*P_o_*, where both drivers were increased simultaneously during the quasi-static computation. Compared to the previous case (d), where *P_h_ ≡ P_o_* remained constant but elastin degraded to a greater extent up to 0.65, the uniform increase in pressure accentuated thickening (λ_*rh*_ = 2.18 at mid thickness) and the change in volume (by mass exchange, mesh (e)-left), but not dilation (λ_*θh*_ = 1.87) or axial retraction (λ_*zh*_ = 0.65) in the most damaged region (*z_o_ = z*_*o*m_, *θ_o_ = π*); it also rendered more uniform the deformation within the crosssection, with λ_*rh*_ = 2.10, λ_*θh*_ = 1.42, and λ_*zh*_ = 0.71 at *θ_o_* = 0. Mechanobiologically equilibrated stresses remained bounded throughout the arterial wall again, reaching mean values *σ_θθh_* = 248 kPa and *σ_zzh_* = 226 kPa at the central cross section.

All FE simulations – even when non-uniform along radial, circumferential, or axial directions and when computed for marked local degradation of elastin (b to d) combined with simultaneous increases in pressure (e) – were computed over 10 incremental global steps during Stage II, with asymptotically quadratic rates of convergence and CPU times (~ 10 min) shown in Table 3. Hence, the implementation is robust and efficient for computing fully resolved G&R responses, with possible numerical issues encountered only when locally approaching mechanobiological static instabilities or excessively distorting the FE mesh, as expected. Indeed, given the computational efficiency, we considered full arterial segment geometries in all of the analyses, with up to 38400 elements, even though by obvious symmetry considerations partial sectors of the specimen could have been analyzed equivalently. Importantly, a recent FE implementation of the original heredity-integral constrained mixture G&R formulation reported 30 hours, on average, to complete a simulation for a 2° cylindrical sector of an axisymmetric fusiform aneurysm meshed with 1200 elements [36]. Thus, the computational time reduces drastically when one employs a rate-independent counterpart of the full formulation, which, with ~ 30 times more elements, and running ~ 150 times faster, roughly represents a 3-fold enhancement in computational efficiency. Nevertheless, the present formulation is not designed to replace the more general heredity-integral formulation in all cases. For example, the present implementation cannot compute transient G&R responses to step changes in blood pressure, flow, or axial stretch, or predict mechanobiological dynamic instabilities, for which the full formulation is yet needed [8, 10, 36].

**Table 3:** Asymptotically quadratic rates of convergence for residual force (left column) and energy (right column) during a typical global time step for the simulation of a hypertensive asymmetric fusiform aneurysm, with an asymmetric degradation of elastin and uniform increase in pressure, shown in Fig. 8 (mesh e), as well as total elapsed CPU time (1 +10 time steps) using a single processor. The FE model comprised *N_r_* × *N_θ_* × *N_z_* = 4 × 160 × 60 = 38400 displacement-based hexahedral elements.

|  | force | energy |
| --- | --- | --- |
| Iteration 0 | 1.2E+00 | 8.7E-01 |
| Iteration 1 | 1.1E-01 | 3.1E-02 |
| Iteration 2 | 2.5E-03 | 3.5E-06 |
| Iteration 3 | 5.5E-06 | 1.0E-10 |
| Iteration 4 | 2.1E-11 | 2.2E-19 |
| Time (1+10 steps) | 11 min 20 s |  |

### 5.4. A parametric study

In the previous examples, we constrained the collagen fibres to deposit along referential orientations 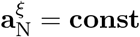 with prestretches 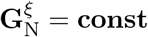. This modeling assumption may be appropriate for abdominal aortic aneurysms (AAAs) [44]. Nonetheless, collagen fiber orientation appears generally orthotropic and widely dispersed at the most dilated sites in AAAs, for example, with a circumferential alignment of collagen fibers observed in some cases [45]. Given the computational efficiency of the present approach, we parametrically test possible in-plane anisotropic effects on the final shape of developed aneurysms by prescribing a gradual re-orientation for the angle *α*_0_ at which symmetric diagonal fibers are deposited during mechanobiologically equilibrated evolutions.

Figure 9 shows contour maps for circumferential (first and third rows of panels) and axial (second and fourth rows) G&R responses of six different aneurysms that result from respective axisymmetric or asymmetric losses of elastic fiber integrity (up to 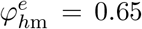) combined with either an alignment *α*_0*h*_ = *α*_0*o*_ 30° that remains constant (left column of panels; cf. meshes (b) and (d) in Figure 8), increases modestly up to *α*_0*h*_ = (4/3)*α*_0*o*_ ≈ 40° (central column), or reorients dramatically up to *α*_0*h*_ = (5/3)*α*_0*o*_ ≈ 50° (right column). In general, a constant alignment for diagonal collagen fibers causes an overall axial retraction of the central region of the aneurysms that induces an expansion of the non-aneurysmal lateral regions (as a result of the axially fixed ends, cf. also meshes (c) in Figure 8 for fixed axial tractions); the modest reorientation causes no overall lengthening of the central region or at the ends; the severe reorientation, in contrast, causes an overall axial expansion of the central region and resulting retraction of the regions next to the fixed ends (as observed in, for example, [40] using a bilayered membrane model with layer-specific fiber orientations).

**Figure 9:**
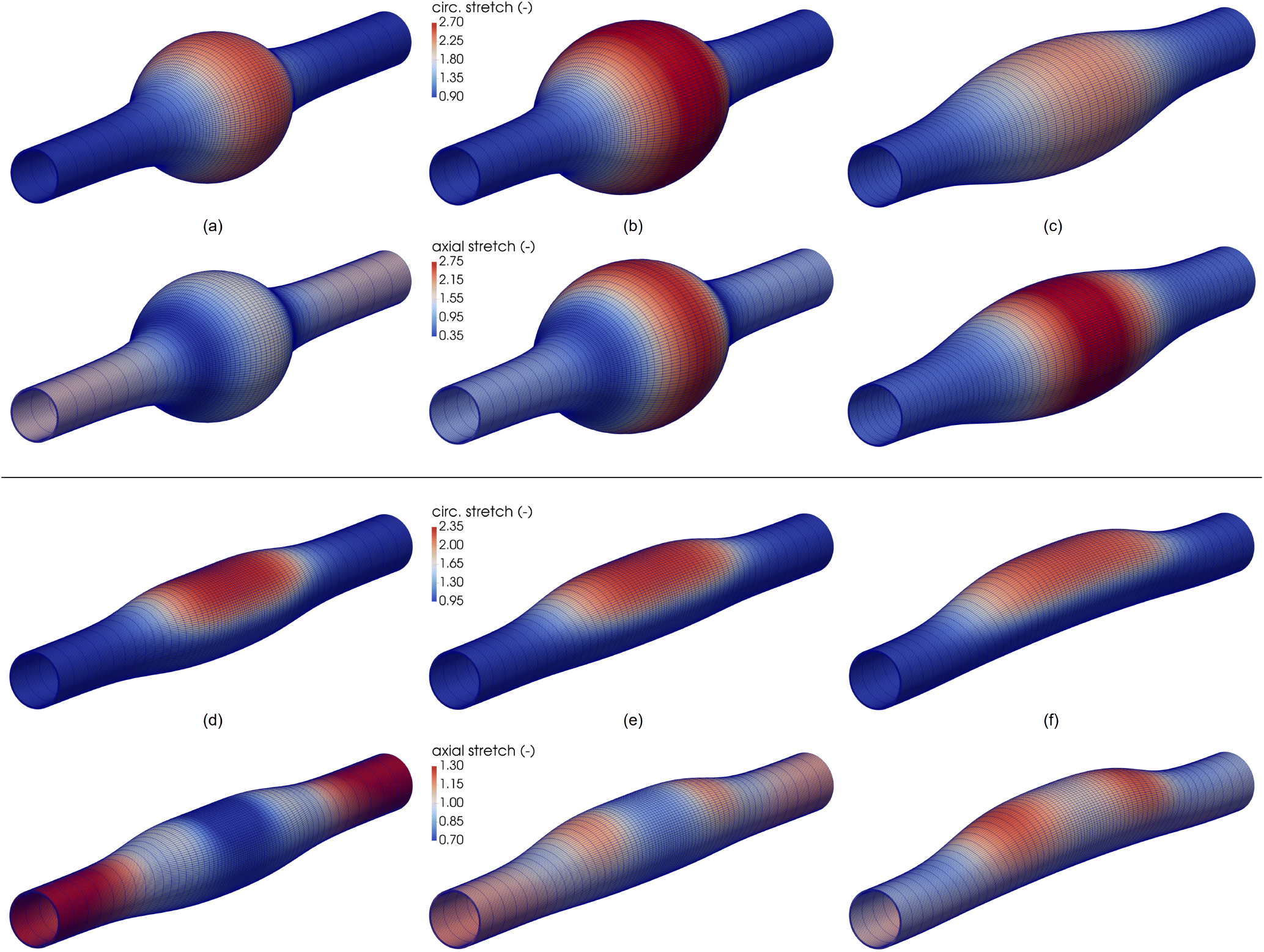
Mechanobiologically equilibrated states for an arterial segment under normotensive conditions (*P_h_* = *P_o_*) with axisymmetrically (a, b, c) or asymmetrically (d, e, f) prescribed elastin damage (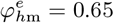 for all cases) in combination with gradually increased orientations of diagonal collagen fibers towards the circumferential direction as (a, d) *α*_0*h*_ = *α*_0*h*_ ≈ 30°, (b, e) *α*_0*h*_ = (4/3) *α*_0*o*_ ≈ 40°, and (c, f) *α*_0*h*_ = (5/3) *α*_0*o*_ ≈ 50°. Shown are deformed meshes and contour plots of circumferential (first and third rows) and axial (second and fourth rows) stretches with respect to an initially straight arterial segment under homeostatic conditions (see mesh (a) in Figure 8). Additional parameters that define the spatially distributed material insults and properties for the axisymmetric or asymmetric aneurysms are the same as described for simulations (b) or (d) in Figure 8. Note the increase in local axial growth within the damaged regions for an increased reorientation of diagonal fibers towards the circumferential direction for both axisymmetric and asymmetric aneurysms (from left to right).

Interestingly, the more pronounced the axial growth during the formation of an asymmetric aneurysm, the more tortuous its luminal centerline, as commonly seen with AAAs, which appears to mark the development of axial-growth-induced tortuosity, that is, an axial-growth-induced lengthening of the aneurysm between fixed ends in the presence of axially and circumferentially geometric and material asymmetries (i.e., in the presence of an asymmetric aneurysm itself).

## 6. Discussion

Many have simulated complex 3D biological growth in diverse soft tissues using a finite (kinematic) growth framework [2, 4, 5, 46–48]. Advantages of this approach include its computational tractability, enabling implementations in existing finite element solvers. A prior disadvantage, however, has been the lack of consideration of the different material properties and rates of turnover of the different constituents that make up native and tissue engineered soft tissues.

In this paper, we presented a new constrained mixture formulation that enables computational tractability while retaining the biologically important characteristic of individual constituent properties, including rates of production and removal. This efficiency was achieved by using a mechanobiologically equilibrated framework, which avoids the heredity integral basis of a full constrained mixture model while providing precise information on the long-term “relaxed” solution [7]. This approach is also useful when time scales associated with G&R are shorter than those associated with perturbations in mechanical or chemical stimuli to which the cells respond [10]. Illustrative solutions for idealized murine arteries subjected to perturbations in blood pressure and flow as well as changes in axial loading or degradation of elastin (Figs. 4 to 7) demonstrated near exact correspondence between the current and prior constrained mixture models. Importantly, the present formulation is also easily implemented within available finite element solvers, though for non-symmetric tangent stiffness matrices. Results were presented for a modified open source code, FEBio, but preliminary simulations showed similar results using ADINA. The associated solutions revealed quadratic convergence with computational efficiency comparable to that for a nonlinear hyperelastic solution, though here for the simultaneous solution of mechanical and mechanobiological equilibrium at load steps that capture evolving geometries, compositions, and properties of interest.

Enlargement of a fusiform aneurysm was simulated easily (Fig. 8), with results similar to previous studies [35, 40, 41, 49], though revealing an important new mechanobiological finding. Not all past simulations of aneurysmal G&R have included both pressure-induced wall stress and flow-induced shear stress in their “stimulus functions” for mass production, and in some cases uncontrollable G&R was noted at the non-aneurysmal ends. The present results demonstrate that flow-induced shear stress-mediated matrix turnover contributes to control the biaxial G&R near the non-aneurysmal ends. This finding is biologically sensible for the endothelial cells should be mechanobiologically responsive, and so too the underlying smooth muscle cells, in these non-aneurysmal end regions. In contrast, endothelial dysfunction could manifest within the enlarged aneurysmal region, where disturbed flows exist and smooth muscle cell drop-out is prevalent. These model predictions during quasi-static mechanobiologically equilibrated evolutions, complemented with others in [36] that show a dynamic stabilization of the expansion of abdominal aortic aneurysms afforded by similar effects of flow-induced shear stresses [8], thus motivate the need for new regional-specific assessments of cell functionality in these lesions.

For simplicity in presenting this new computational G&R framework, we did not consider specific criteria for elastin degradation (e.g., induced by low wall shear stress in fluid-solid growth computational frameworks [50–52]) or damage (e.g., induced by excessive stretching [49]), which was prescribed herein as the stimulation driver for the quasi-static computations of aneurysmal enlargement. Simulated effects of varying deposition stretches, rates of turnover, and material stiffness of collagen have been previously investigated parametrically and proven to play fundamental roles in the natural history of AAAs [53, 54]. We explored effects of the degree of wall heterogeneity and anisotropy on the resulting G&R response in terms of varying orientations of the newly incorporated collagen fibers within the aneurysmal wall (Fig. 9). Consistent with previous studies that predicted local axial expansions influenced by marked axial off-loadings and noted a propensity for the aneurysm to buckle [40, 54], our results revealed that increasing ratios of axial to circumferential growth emerged for a prescribed marked loss of local elastic fiber integrity (up to 65%) in combination with a gradual reorientation of diagonal fibers of collagen towards the circumferential direction, which led to increasing out-of-plane deformations of the aortic centerline for the asymmetric aneurysms. Collectively, these results highlight the importance of considering anisotropic G&R when modeling biological tissues [55, 56] and, especially, of quantifying not only circumferential growth responses (i.e., maximum diameter) but also axial growth responses (i.e., overall lengthening), along with their expansion rates, when assessing the potential of aneurysms to rupture [57, 58]. In this regard, the present approach need not prescribe a predefined directionality for the G&R or associated deformation tensors; rather, case-specific G&R responses emerge from the simultaneous solution of mechanical (Eq. (7)_2_) and mechanobiological (Eq. (6)_1_) equilibrium equations complemented by mechanobiologically-inspired constitutive relations for constituent stresses (Section 2.5) and stimulus functions (Section 2.6) subject to case-specific boundary conditions (see also [56]).

Although full, heredity-based constrained mixture models (e.g., [12, 15, 31, 32]) will continue to be needed to study particular physiological and pathophysiological situations, the present formulation appears to apply in many cases, provided that the loading or structural perturbations remain static or are slow enough relative to the G&R process. We submit, therefore, that this easily implemented, fast, rate-independent finite element approach to modeling soft tissue G&R will enable previously impractical parametric studies that can be used to evaluate novel mechanobiological hypotheses, which in turn can better focus cost- and time-intensive experimental studies for a host of soft tissues [11].

There is, of course, further need to understand better the actual mechanical stimuli that the cells sense and respond to, noting that different cells may respond to different magnitudes or even different stimuli. We considered *σ_υo_* and *τ_wo_* as homeostatic targets, which yielded numerical responses that reflect experimental observations. Much more remains to be discovered, and then modeled, in soft tissue mechanobiology.

## Acknowledgments

This work was supported, in part, by grants from the NIH (R01 HL128602, P01 HL134605, U01 HL142518) and DoD (W81 XWH1810518).

## Appendix A. Deformation gradients during general G&R responses

Within a general constrained mixture theory, the deformation gradient for a constituent *α* deposited at time *τ* within extant matrix, and that survives at current G&R time *s* ≥ *τ*, reads [12]

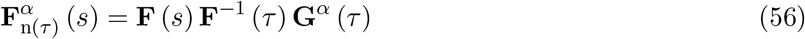

which maps line elements (fibers) from an evolving natural configuration 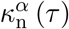 to the current configuration of the mixture *κ*(*s*), with **G**^*α*^ (*τ*) = (**G**^*α*^ (*τ*))^T^, and det **G**^*α*^ (*τ*) = 1. At deposition time *τ*,

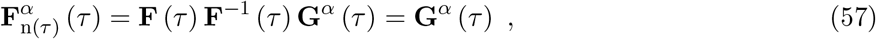

thus the natural configuration 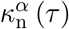 evolves with the current configuration of the soft tissue at *τ*. It is for this reason that newly produced collagen families are often assumed to be incorporated within the wall, at a homeostatic stretch, along unit vectors relative to *spatial* directions (i.e., eigenvectors) of the Cauchy stress tensor in the loaded configuration at time of deposition [12, 14, 59]. Consequently, 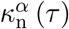 is subject to possible rigid-body motions superimposed on the current configuration at *τ*. In other words, the magnitude of **G**^*α*^ (e.g., its eigenvalues) might be assumed to remain constant over time, but not so its spatial orientation (i.e., its eigenvectors).

To develop formulations insensitive to rigid body motions, it is often convenient to define equivalent variables in a referential (Lagrangian) description. Hence, upon considering the right polar decomposition of the deformation gradient

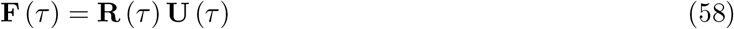

and, because of the inherent link between 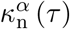 and the corresponding current configuration of the mixture *κ*(*τ*), we can define a (yet symmetric, volume-preserving) prestretch tensor rotated by **R**^T^ (*τ*) as

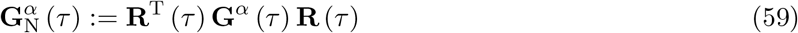

which is determined relative to an evolving natural configuration 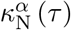 that is rotated with respect to 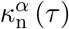 but remains unrotated with respect to the reference configuration of the mixture *κ*(0) = *κ_o_*. This motivates the introduction of a two-point linear transformation defined at the constituent level as

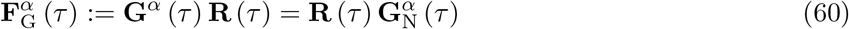

which, importantly, allows one to interpret (the original) **G**^*α*^ and (the herein introduced) 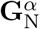 as respective *left* and *right* prestretch tensors defined with respect to the Lagrangian-type natural configuration 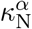. Therefore, we can also define (cf. Eq. (56))

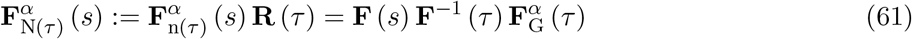

which maps line elements (fibers) from 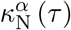 to the current configuration of the mixture *κ*(*s*). Hence, at deposition time *τ* (cf. Eq. (57))

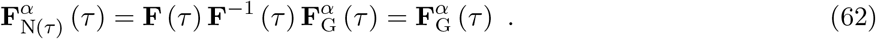

In this way, one can conveniently remove possible sources of non-objectivity by considering the rotated (right) deposition stretch tensor 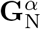. Here, we posit, for example, that both its magnitude and orientation remain constant

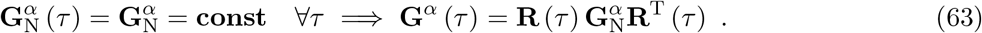

In cases that rotations are absent, as, for example, for axisymmetrically loaded arteries treated as thin-walled [7, 8] or thick-walled [14] cylinders, this assumption yields 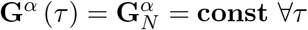, as assumed therein.

For mechanobiologically equilibrated states, when **F** (*τ*) = **F** (*s*) =: **F**_*h*_, we have, in general

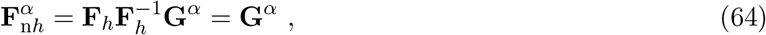

and

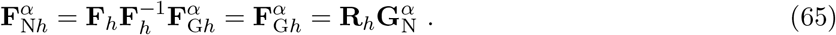

The latter decomposition, expressed in terms of **R**_h_ ≠ **const** and 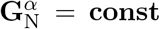, proves useful when defining mechanobiologically equilibrated (rotated) Cauchy stresses subject to objective requirements. In this regard, note that an arbitrary rigid-body rotation **Q** ∈ orth^+^ superimposed on 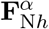 yields

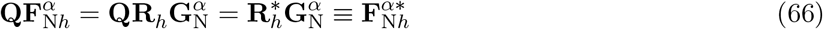

with the superimposed rotation absorbed by 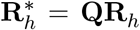, and 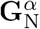 remaining unaltered (i.e., constant, defined in 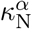). In contrast, because 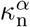 evolves with the current configuration of the mixture, an arbitrary rotation superimposed on 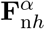 implies a rotation on 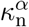 and, hence, enforces a rotation of **G**^*α*^ by means of

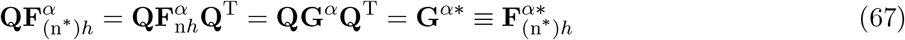

with **G**^*α*\*^ = **QG**^*α*^**Q**^T^ defined in a rotated natural configuration 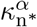, and **Q**^T^ ≡ **Q**^−1^ rotating line elements from 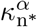 to 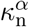 accordingly. Again, note that 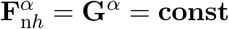 for cylindrical geometries [7, 8].

## Appendix B. Stresses during transient hyperelastic responses

The energy *W*_R_(**C**_*h*_) in Eq. (23) gives that stored by the mixture as a consequence of its pre-stretched in vivo state. To determine associated pre-stresses by differentiation of *W*_R_ under the assumption that the bulk modulus of the tissue far exceeds its shear modulus (i.e., volume may change with growth but not loading over short periods), one can consider arbitrary isochoric (elastic) deformations superimposed to the equilibrated current configuration of the mixture *κ_h_*. Particularization of these expressions to **F** ≡ **F**_*h*_ provides mechanobiologically equilibrated stresses.

Consider the combined gradient 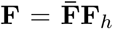, with **F**_*h*_ fixed and 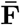 superimposed. From 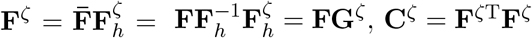, **C**^*ζ*^ = **F**^*ζ*T^**F**^*ζ*^ yields, with **C** = **F**^T^**F**,

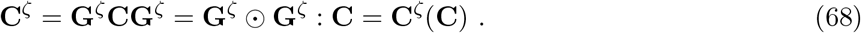

Similarly, from 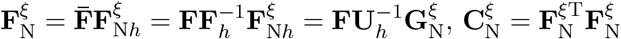 yields

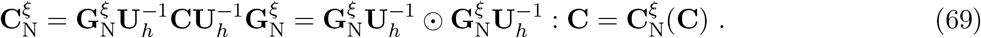

Since 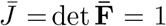, then 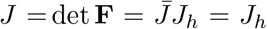, and the strain energy for the mixture, particularized to a homeostatic state in Eq. (23), generalizes to

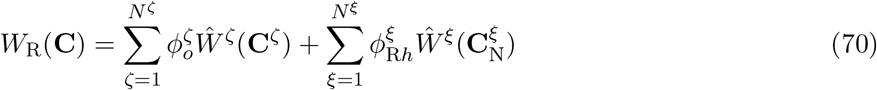

with 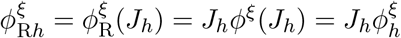 remaining constant.

The stress power per unit reference volume in *κ_o_* is, for this intermittent hyperelastic response, 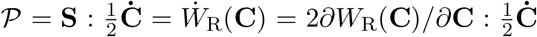, with **S** the second Piola–Kirchhoff stress for the mixture, which yields

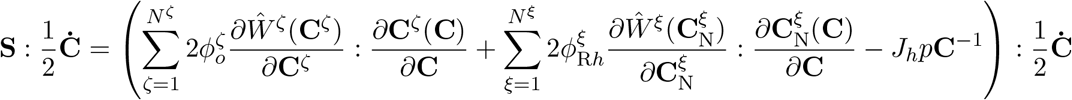

where we accounted for dependencies in Eqs. (68) and (69) and added the last term consistent with the kinematic constraint 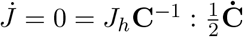, with *p* the associated Lagrange multiplier. Then, 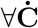 subject to 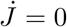, we obtain

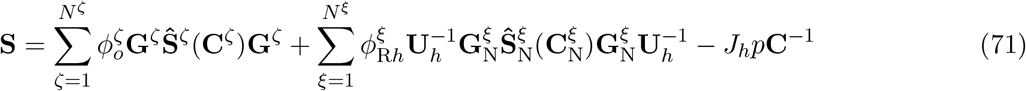

where we defined second Piola–Kirchhoff stresses at the constituent level for both type of constituents

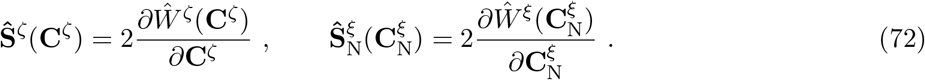

Note that Eqs. (70) and (71), obtained for hyperelastic (isochoric) responses with respect to an evolved, grown and remodeled, state *κ_h_*, constitute a generalization of both the mass averaged rule-of-mixture strain energy and stresses in Ref. [30], obtained for hyperelastic (isochoric) responses with respect to an original homeostatic state *κ_o_*, for which *W*_R_ ≡ *W*, 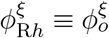, *J_h_* ≡ *J_o_* = 1, and **U**_*h*_ ≡ **U**_*o*_ = **I**, namely

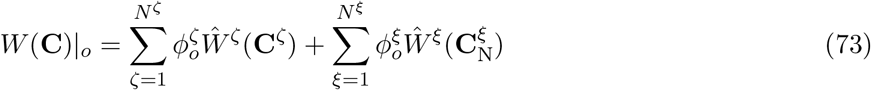

and

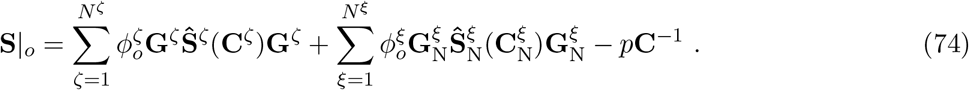

## Appendix C. A stimulus function for arterial G&R

In previous works, we have analyzed G&R of prototypical cylindrical arteries with stimulus functions for mass production

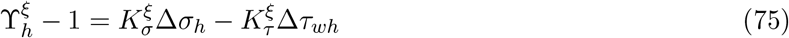

where

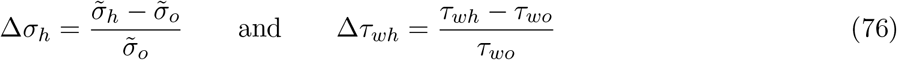

with 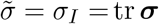 the first principal invariant of the mean wall stress and *τ_w_* = 4*μQ*/*πα*^3^ a measure of the wall shear stress over the endothelium for a fully developed Newtonian flow through a long cylindrical sector, with *μ* the viscosity of blood, *Q* the volumetric flow rate, and *a* the current luminal radius. Note that, while we can still employ 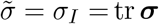 in the present general framework, with ***σ*** now defined pointwise in Eq. (32), we cannot compute local values for *τ_w_* for complex geometries, for which one would need to incorporate computations for the blood flow to account for fluid-solid interactions, namely perform fluid-solid-growth cardiovascular simulations [60] particularized in the present case to quasi-steady flows.

### Intramural and wall shear stress stimuli: general case

For quasi-steady-state fluid-solid-growth formulations modeled with the present framework, fluid and solid computations interact through the mechanobiologically equilibrium condition 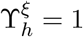. From Eq. (75), considering also proportional out-of-equilibrium stimulus functions **ϒ** − 1 for smooth muscle and collagen, such that 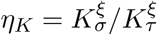 [7], we arrive at

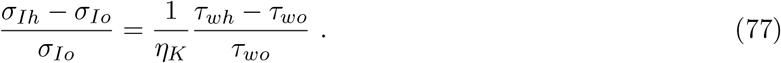

From Eq. (32) we obtain 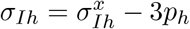, which leads to

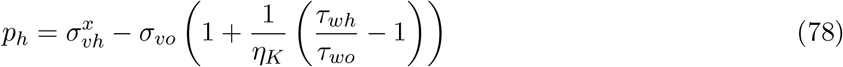

with the volumetric stresses 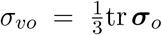 (known from the original homeostatic, loaded state) and 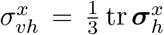. Importantly, note that Eq. (78) couples solid stress computations, through Eq. (32), with blood flow computations, through the wall shear stress measure *τ_wh_*. In particular, the volumetric Cauchy stress for the mixture 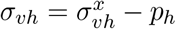 in Eq. (32) reads

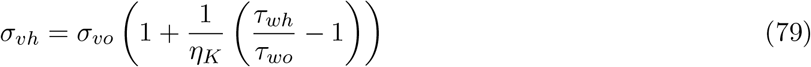

where, consistent with the considered stimulus function, the flow-induced wall shear stress modulates the intramural volumetric stress.

### Intramural and wall shear stress stimuli: cylindrical arterial segments

Consider a long cylindrical artery that remains cylindrical over G&R time scales, for which the wall shear stress expression *τ_w_* = 4*μQ*/*πα*^3^ applies. Yet, this analytical expression for *τ_w_* cannot be assessed locally in our arterial wall model because of the presence of the luminal radius *a*, which represents a geometrical outcome of the boundary value problem (i.e., the computed radial coordinate of the inner surface). However, we can relate the global variable *a* with variables defined locally within a cylindrical arterial wall, at a given current radial coordinate *r*, satisfying *α* ≤ *r* ≤ *r*_out_, with *r*_out_ the current outer radius, as

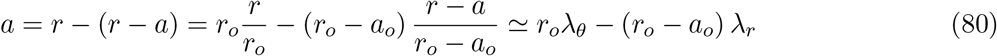

where *r_o_* is the original radial coordinate, *r*/*r_o_* = λ_*θ*_ is the (exact) local circumferential stretch computed from **C** = **F**^T^**F** as 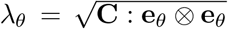, and (*r* − *a*) / (*r_o_* − *a_o_*) ≃ λ_*r*_ is an (approximated) local radial stretch computed as 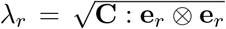, with **e**_*θ*_ and **e**_*r*_ unit vectors along circumferential and radial directions. With 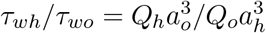, Eq. (78) becomes

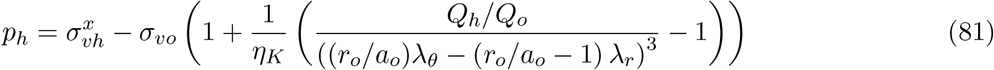

which, compared to Eq. (36), requires additional straightforward tangent contributions in Eq. (48) expressed in terms of the derivatives

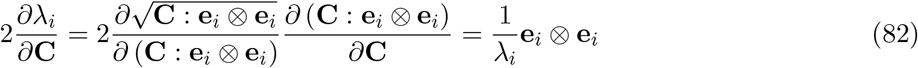

for both *i* = *r, θ*, and that we used to illustrate qualitative results for arterial G&R in examples above.

## Appendix D. Derivation of tangent moduli contribution 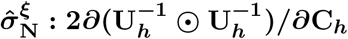

We derive here the expression for a fourth-order tensor 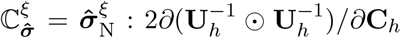 based on spectral decompositions for the rate of change of the different tensors involved [61, 62]. For notation simplicity, we disregard subscript *h* in deformation tensors. The spectral decomposition of the right Cauchy–Green tensor **C** reads

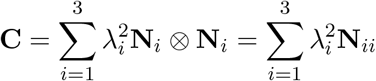

where λ_*i*_ are stretches and **N**_*i*_ Lagrangian strain eigenvectors, and we define eigentensors **N**_*ab…c*_ := **N**_*a*_ ⊗ **N**_*b*_ ⊗ … ⊗ **N**_*c*_. The material time derivative of 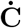 yields

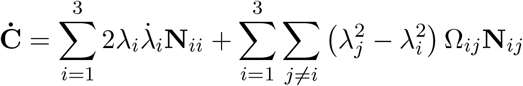

where 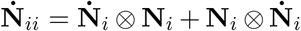 and Ω_*ij*_ = –Ω_*ji*_ are components of the angular velocity (skew) tensor of Lagrangian eigenvectors **Ω** expressed in that same basis, such that

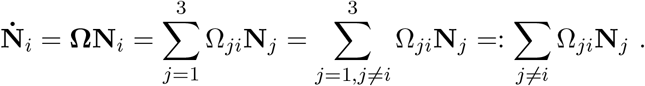

Consider now the spectral decomposition of the inverse of the right stretch tensor

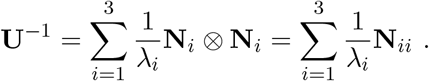

The spectral decomposition of the fourth-order tensor **U**^−1^ ⊙ **U**^−1^ is

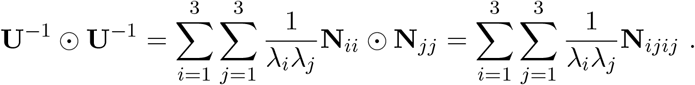

Its material time derivative, proceeding similarly as we did for 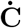, and after few algebraic (index) manipulations, reads

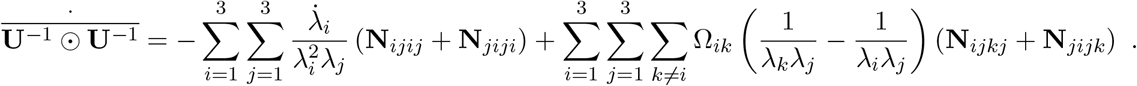

Since 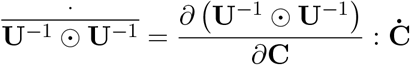, by comparing expressions for 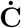 and 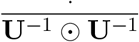, one identifies the sixth-order tensor

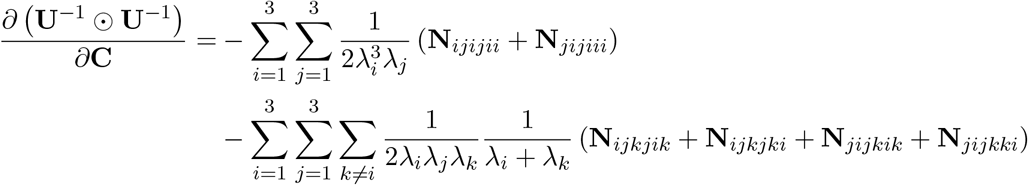

which could be implemented, as is, in finite element codes to be subsequently doubly contracted, numerically, with 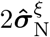 to give 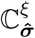. More efficiently, however, we perform the double contraction analytically, and the result is implemented directly as a fourth-order tensor in our user-defined material subroutine. Consider then the spectral decomposition for the rotated stress tensor 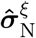 in terms of the eigentensors **N**_*ij*_, which are not coaxial, in general, thereby

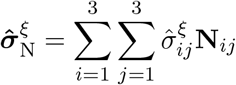

from which 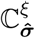 is obtained as

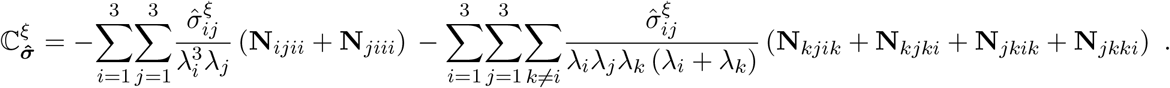

Importantly, note that the fourth-order tensor 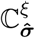 preserves minor symmetries (*abcd*) ↔ (*bacd*) ↔ (*abdc*), but (generally) not the major symmetry (*abcd*) ↔ (*cdab*).

## Appendix E. Specialization: an equivalent thin-walled artery

We reformulate here the main algebraic equations derived in [7] to show their consistency with the general boundary value formulation above.

### Formulation in [7]

We solve a system of nonlinear algebraic equations formed by mechanobiological equilibrium ϒ_*h*_ = 1, the constraint 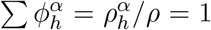, and mechanical equilibrium along both (in-plane) circumferential and axial directions of a cylindrical artery, namely

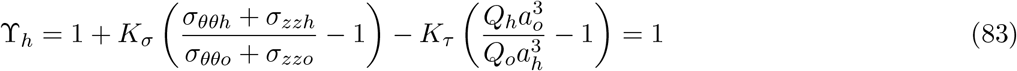

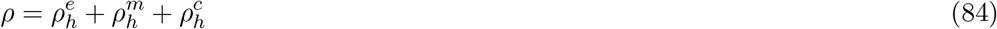

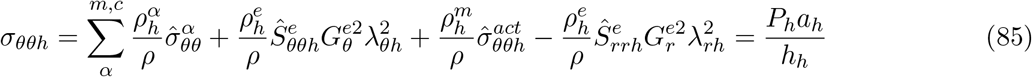

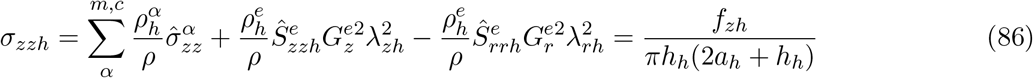

where the primary unknowns are *a_h_, h_h_*, 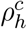, and *f_zh_*. In particular, the evolved homeostatic Lagrange multiplier *p_h_* (not shown) is obtained from the (not shown) radial equilibrium equation and associated boundary condition (*σ_rrh_* = 0), which enables substitution of *p_h_* in expressions for in-plane stresses, hence, representing a specific resolution procedure for a cylindrical artery.

### An equivalent formulation aimed for finite element implementation

One can alternatively solve an equivalent system of nonlinear algebraic equations formed by mechanobiological equilibrium ϒ***h*** = 1 plus mechanical equilibrium along the (out-of- and in-plane) radial, circumferential, and axial directions, namely

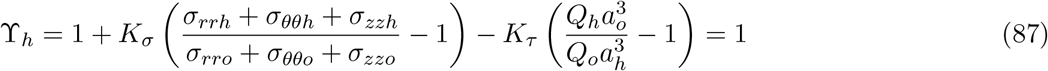

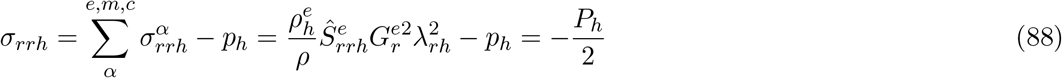

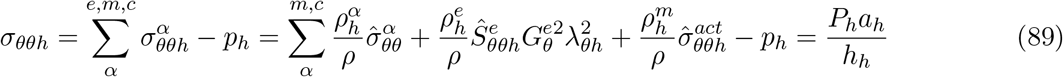

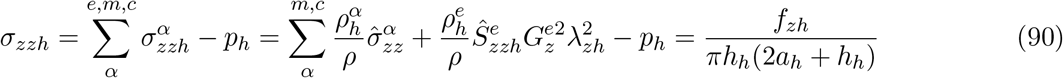

where the primary unknowns are *a_h_, h_h_, p_h_*, and *f_zh_*, noting in Eqs. (87) and (88) additionally the consideration of a mean radial stress – *P_h_*/2 for more accurate comparisons with FE analyses. In particular, the evolved homeostatic Lagrange multiplier *p_h_* can be obtained in this case from the mechanobiological equilibrium equation (recall Eq. (33)), which enables substitution of *p_h_* in expressions for out-of- and in-plane stresses (recall Eq. (32)), hence, representing a generalized resolution procedure for a cylindrical artery consistent with the general formulation derived above.

